# Interpreting the molecular mechanisms of disease variants in human transmembrane proteins

**DOI:** 10.1101/2022.07.12.499731

**Authors:** Johanna Katarina Sofie Tiemann, Henrike Zschach, Kresten Lindorff-Larsen, Amelie Stein

**Author notes:** Correspondence (KL-L), (AS). Center for Health Data Science, Faculty of Health and Medical Sciences, University of Copenhagen, DK-2200 Copenhagen N., Denmark.

## Abstract

Next-generation sequencing of human genomes reveals millions of missense variants, some of which may lead to loss of protein function and ultimately disease. We here investigate missense variants in membrane proteins — key drivers in cell signaling and recognition. We find enrichment of pathogenic variants in the transmembrane region across 19,000 functionally classified variants in human membrane proteins. To accurately predict variant consequences, one fundamentally needs to understand the reasons for pathogenicity. A key mechanism underlying pathogenicity in missense variants of soluble proteins has been shown to be loss of stability. Membrane proteins though are widely understudied. We here interpret for the first time on a larger scale variant effects by performing structure-based estimations of changes in thermodynamic stability under the usage of a membrane-specific force-field and evolutionary conservation analyses of 15 transmembrane proteins. We find evidence for loss of stability being the cause of pathogenicity in more than half of the pathogenic variants, indicating that this is a driving factor also in membrane-protein-associated diseases. Our findings show how computational tools aid in gaining mechanistic insights into variant consequences for membrane proteins. To enable broader analyses of disease-related and population variants, we include variant mappings for the entire human proteome.

**SIGNIFICANCE:** Genome sequencing is revealing thousands of variants in each individual, some of which may increase disease risks. In soluble proteins, stability calculations have successfully been used to identify variants that are likely pathogenic due to loss of protein stability and subsequent degradation. This knowledge opens up potential treatment avenues. Membrane proteins form about 25% of the human proteome and are key to cellular function, however calculations for disease-associated variants have not systematically been tested on them. Here we present a new protocol for stability calculations on membrane proteins under the usage of a membrane specific force-field and its proof-of-principle application on 15 proteins with disease-associated variants. We integrate stability calculations with evolutionary sequence analysis, allowing us to separate variants where loss of stability is the most likely mechanism from those where other protein properties such as ligand binding are affected.

## INTRODUCTION

Proteins carry out the majority of functions in a cell. While most proteins are robust to some sequence changes (1), other single amino acid variants may render them non-functional. For nuclear and cytosolic proteins, we and others have shown that the molecular reason underlying loss of function for human pathogenic variants is often loss of protein stability (2–10). Proteins affected by such destabilizing variants are recognised by the cellular protein quality control system, leading to degradation and hence low levels that cause a loss-of-function phenotype (11). For soluble proteins, structure-based calculations of stability changes upon mutation (ΔΔG) (12) correlate with experimental stability (13–16) as well as high-throughput abundance measurements (17, 18), allowing us to annotate variants accordingly (19). The loss of stability induced by such variants often leads to cellular protein degradation. This mechanistic link to degradation is not only interesting from a biophysical perspective but can also lead to development of treatments that rescue the variant from degradation (20, 21).

Twenty-three percent of sequences in the human proteome encode membrane proteins, including channels, transporters, enzymes, and receptors such as G-protein coupled receptors (GPCRs) (22). Located at the junction between two compartments and often exposed to small molecules in the bloodstream, membrane proteins are key in cell signaling and recognition, as well as major drug targets (23, 24). Variants in membrane proteins are associated with a number of diseases, including for example cystic fibrosis, Parkinson’s, Alzheimer’s and atherosclerosis (25–28).

Studying membrane proteins experimentally or computationally is challenging, as the proteins need to be considered in context of the lipid membrane (29). Furthermore, while many soluble proteins can unfold and refold reversibly, the processes of synthesis, folding, and assembly are intrinsically linked for membrane proteins (30, 31). In particular, denaturants can perturb properties of the membrane (or its mimetics) when thermodynamic stability measurements are performed in (near) native conditions. More recent techniques such as steric trapping or label-free differential scanning fluorimetry aim to avoid those drawbacks but cannot be applied in a high-throughput manner (32, 33). Therefore, large-scale and easily accessible experimental data for benchmarking computational tools are sparse. Despite recent methodological advances (34), computational methods for membrane proteins are not as developed as those for soluble proteins. Furthermore, the diverse experimental studies measure different levels of unfolding, which further challenges computational method development. Thus, the application of computational analyses for examining a potential correlation between protein stability, cellular abundance, and function analogous to that for soluble proteins may be particularly challenging for membrane proteins.

Building on recent force field developments that make computational analysis of membrane proteins more realistic (35), we here set out to assess whether calculations of the change in folding free energy can be used to identify the subset of pathogenic variants that are likely caused by loss of stability. In particular, we calculate the change in folding energy between a wild-type protein and a protein variant (ΔΔG = ΔG_MUT_ - ΔG_WT_), where low ΔΔG correspond to substitutions that—in light of stability—appear well-tolerated, and high ΔΔG for variants that destabilize the protein structure. Of note, the levels of unfolding or destabilization *in vivo* do not necessarily have to lead to complete protein unfolding. Partial unfolding may be sufficient to trigger recognition by the protein quality control system. We first combined several protein annotation databases to obtain an overview of the number and types of missense variants that are found in membrane proteins. We then analyzed in more detail 15 human membrane proteins for which high-resolution structural data as well as annotations of pathogenic and benign variants were available, and calculated ΔΔG values for them. In addition, we used an evolutionary sequence analysis approach (36) to calculate a value which we term ΔΔE indicating the evolutionary importance of each residue. This and similar approaches have been shown to be useful in detecting detrimental variants and includes both loss of stability variants and variants that lose function due to—for example—catalytic impairment of enzymes or mistrafficking (37–39). Multiple recent works have demonstrated that sequence analysis of conservation is able to capture such non-beneficial variants with high accuracy (37, 40, 41). The mechanistic reasons for why a variant is not tolerated by evolution, whether it be gain or loss of function or other aspects such as loss of stability, are not directly apparent, as is the case for many predictors. In the following, we label ΔΔE with *loss of function* to facilitate reading. In this work, we use it in combination with loss of stability for dissection of underlying causes. The combination of ΔΔG and ΔΔE has proven particularly useful for providing mechanistic insight into loss-of-function variants in soluble proteins (18, 19). We here apply such a combined analysis to gain mechanistic insight into variant consequences in 15 selected membrane proteins.

## METHODS AND MATERIALS

### Collection and processing of clinical, population and structural data

To extract all annotated human membrane proteins, we first obtained all unique proteins (UniProt-ID) of the human proteome (*organism=homo sapiens*) from the UniProt (https://www.uniprot.org/help/api) (42) and EMBL-EBI (https://www.ebi.ac.uk/proteins/api/doc/) database. For each UniProt-ID, we then stored its general and amino acid-based annotations (such as protein domain regions) in UniProt and further selected proteins of the type ‘TRANSMEM’, ‘INTRAMEM’, ‘TOPO_DOM’, or ‘LIPID’. This annotation originates from assignment of structural properties or predictions by TMHMM (https://www.uniprot.org/help/topo_dom). The UniProt-ID of the first transcript is used in the further mapping and analysis.

We then further filtered the UniProt-ID list so that all remaining proteins have at least one ClinVar (43) or gnomAD (44) missense variant. gnomAD data were taken from an in-house database built on exome data from gnomAD v2 and whole-genome data from gnomAD v3 (scripts available at https://github.com/KULL-Centre/PRISM/tree/main/software/make_prism_files, release-tag v0.1.1). The database was generated by first downloading the vcf files (May 2021), selecting exome GRCh38 liftover for v2 and whole-genome files for v3. The vcf files were then annotated with Variant Effect Predictor (VEP) with the GRCh38 release 100 human transcripts set from Ensembl. From the annotated vcfs we established for all protein-level variants, separately in exome and genome data, allele frequencies from the variant allele count as the sum of all DNA variants leading to the same protein-level variant. Clinvar data were obtained by parsing the following file: https://ftp.ncbi.nlm.nih.gov/pub/clinvar/tab_delimited/variant_summary.txt.gz (May 2021) and only admitting entries that have a rating of at least one star, are single nucleotide variants and mapped to GRCh38 (script available at https://github.com/KULL-Centre/PRISM/tree/main/software/make_prism_files, release-tag v0.1.1). The dataset for the entire human proteome is provided at https://sid.erda.dk/sharelink/c3rDfqR8nn, using a UniProt-AC-based directory structure, e.g., files for GTR1 (UniProt AC P11166) can be found in subdirectory prism/P1/11/66/.

Next, we extracted all PDB-IDs from RCSB and PDBe with a matching UniProt-ID reference. As not every PDB-ID for a given UniProt-ID in PDBe could be found in RCSB PDB and vice versa, we searched both databases. We further included phenotype and genomic disease annotations from OMIM via mim2gene (https://omim.org/static/omim/data/mim2gene.txt) and MIM, including the proteins’ chromosome information.

The sequences were then aligned to the UniProt sequence using *pairwise2.align.globalds* (with BLAST defaults) from Biopython (45), a minimal identity of 0.6, and minimal coverage of 0.1 for alignment acceptance. All residues that do not match the UniProt sequence were discarded. The final data contained each protein sequence, its UniProt-ID, the secondary structure prediction by residue, solvent-accessible surface area for each wild-type residue, and UniProt annotations such as transmembrane region, protein modifications, total allele frequency counts from gnomAD, ClinVar significance statements, genomic disease annotations, and associated PDB-IDs.

### Selection of targets used for computational predictions

To find a set of proteins for our computational sequence and structure analyses, we selected all proteins that have at least one benign and one pathogenic ClinVar annotation in an experimentally resolved transmembrane region of the protein. This reduced the number of proteins with gnomAD or ClinVar annotations from 1,504 proteins to 41 proteins. As the Rosetta membrane energy function has been developed and benchmarked on structures resolved by X-ray crystallography, we selected those, reducing the protein set to 16. The selected proteins are listed in table 3 and Supporting Data at https://github.com/KULL-Centre/papers/tree/main/2022/hMP-Xray-Tiemann-et-al.

The structures for each of the chosen proteins have been selected according to their *Structure Selection Score* (StrucSe_score) and the number of variants in total and within the transmembrane region. The StrucSel_score is a combination of method resolution, sequence coverage and identity to the experiment and the wild-type (according to UniProt), including an annotation about inserts, deletions, mismatch, non-observes and modified residues. The script is available at https://github.com/KULL-Centre/PRISM/blob/main/software/scripts/struc_select_sifts.py and the table with the numbers for each of the proteins at https://github.com/KULL-Centre/papers/blob/main/2022/hMP-Xray-Tiemann-et-al/data (*date*-count_hMP_anno_splitPDB_Xray_publish.xlsx).

### Conservation analysis of variant effects

To calculate the effect of a variant in light of evolution, we used the GEMME algorithm (36) as previously described (19): We first construct a multiple sequence alignment (MSA) using the sequence of the first transcript of each proteins UniProt-ID as input to HHBlits (version 2.0.15) (46) with the following settings -e 1e-10 -i 1 -p 40 -b 1 -B 20000 to search UniRef30_2020_03_hhsuite.tar.gz (47–49). The MSA is filtered by keeping only positions present in the target sequence and sequences with less than 50% gaps. We then further follow the GEMME algorithm that predicts the degree of conservation for all 19 substitutions (ΔΔE). We rank-normalized the ΔΔE values for the entire protein to allow comparison with the other proteins in the dataset. ΔΔE ≈ 0 corresponds to well-tolerated substitutions, whereas ΔΔE ≈ 1 corresponds to rare or absent variants. Additionally, we extract the sequence coverage of the MSA for each position.

### Thermodynamic stability predictions

To calculate changes in thermodynamic stability (ΔΔG), we use Rosetta version v2021.31-dev61729-0-gc7009b3115c (GitHub sha1 c7009b3115c22daa9efe2805d9d1ebba08426a54). We implemented an in-house pipeline to perform preparation, relaxation, and ΔΔG calculations of the protein (https://github.com/KULL-Centre/PRISM/tree/main/software/rosetta_ddG_pipeline, release-tag v0.1.1). Preparation includes cleaning of the PDB structure coordinates (hereafter referred to as structure) of ligands and alternative rotamers and chains, superposing of the protein into the membrane plane as well as calculation of the membrane plane, lipid-accessible residues (50) and the solvent-accessible surface area using DSSP (51, 52) (the latter is solely used for analysis purposes).

To utilize the membrane protein mode in Rosetta, two conditions must be met: first, a membrane plane file, containing the residues that are within the membrane, needs to be provided; second, the structure of the protein must be centered and oriented within the membrane, where the membrane thickness follows the z-axis. The membrane plane can be calculated using a membrane-aligned protein structure. Therefore, protein coordinate translation was performed by structural superposition of the protein to its equivalent structure obtained from the *Orientations of Proteins in Membranes (OPM) database* (53), which lies already within those coordinates. (If the chosen PDBid is not present in the OPM database, an alternative structure for the same protein or a close homolog is chosen.) Next, the membrane plane was calculated using Rosetta (54, 55) and the protein structure was relaxed as described in (56). Finally, ΔΔG values for each variant were calculated as the residual energy of the variant minus the energy of the wild type.

We performed a benchmark to identify the best protocol to calculate ΔΔGs for membrane proteins. First, we collected 20 experimentally derived ΔΔG datasets of in total 8 different membrane proteins (Table S3). Then, we implemented three different protocols, namely MP_repack, MP_flex_relax_ddG, and ‘cart_prot’, inspired by work on soluble proteins (12) and previously published work on membrane proteins (35). MP_repack operates in *torsion space* and performs a simple repacking of the side chains within a defined radius after mutagenesis (following the protocol mentioned in (35)). MP_flex_relax_ddG is inspired by (12) and allows more flexibility to accommodate the variant by allowing backbone relaxation of the variant and its sequential neighbors, in addition to repacking of side chains within a defined radius. cart_prot follows the same protocol as MP_flex_relax_ddG but is executed in *cartesian space*. For all protocols, we used the membrane protein score function ‘franklin2019’ (35) that performs comparable to older membrane scoring functions as recently evaluated in (56). Finally, we selected cart_prot as the computed values gave the best correlation with the experimental data (0.46), and additionally, the computed values have a high reproducibility, indicated by the low standard deviation for replicates (Fig. S3). As mentioned in the Limitations section, the correlation of independent experimental studies on the same protein and the same variants (57, 58) is 0.65.

### Enrichment of benign variant counts by gnomAD allele frequency

To evaluate the value of our computational methods to predict variants to be benign or pathogenic using ROC analysis, we aimed for a large number of benign and pathogenic variants. In our target proteins, we have 324 pathogenic but only 122 benign variants. We aimed to supplement benign variants with variants from gnomAD. Therefore, we performed a ROC analysis of the gnomAD allele frequency on the 10,260 benign and 2,360 pathogenic ClinVar variants in the human membrane proteome that also have a gnomAD allele frequency (Fig. 1C) and obtained an AUC of 0.96 (Fig. S1). This analysis enables calculating a cutoff to separate benign from pathogenic variants using gnomAD allele frequency via the highest Youden Index (59)). We thereby obtain a cutoff of 9.9 · 10^-5^ (Fig. 1B). We define *group B* variants as the union of those that are defined by ClinVar as benign and those variants that have an allele frequency > 9.9 · 10^-5^ and are not pathogenic (in ClinVar). Consequently, we call pathogenic variants *group A*. This results for our target proteins in 324 *group A* and 283 *group B* (benign and/or non-rare) variants across 16 proteins.

**Figure 1:**
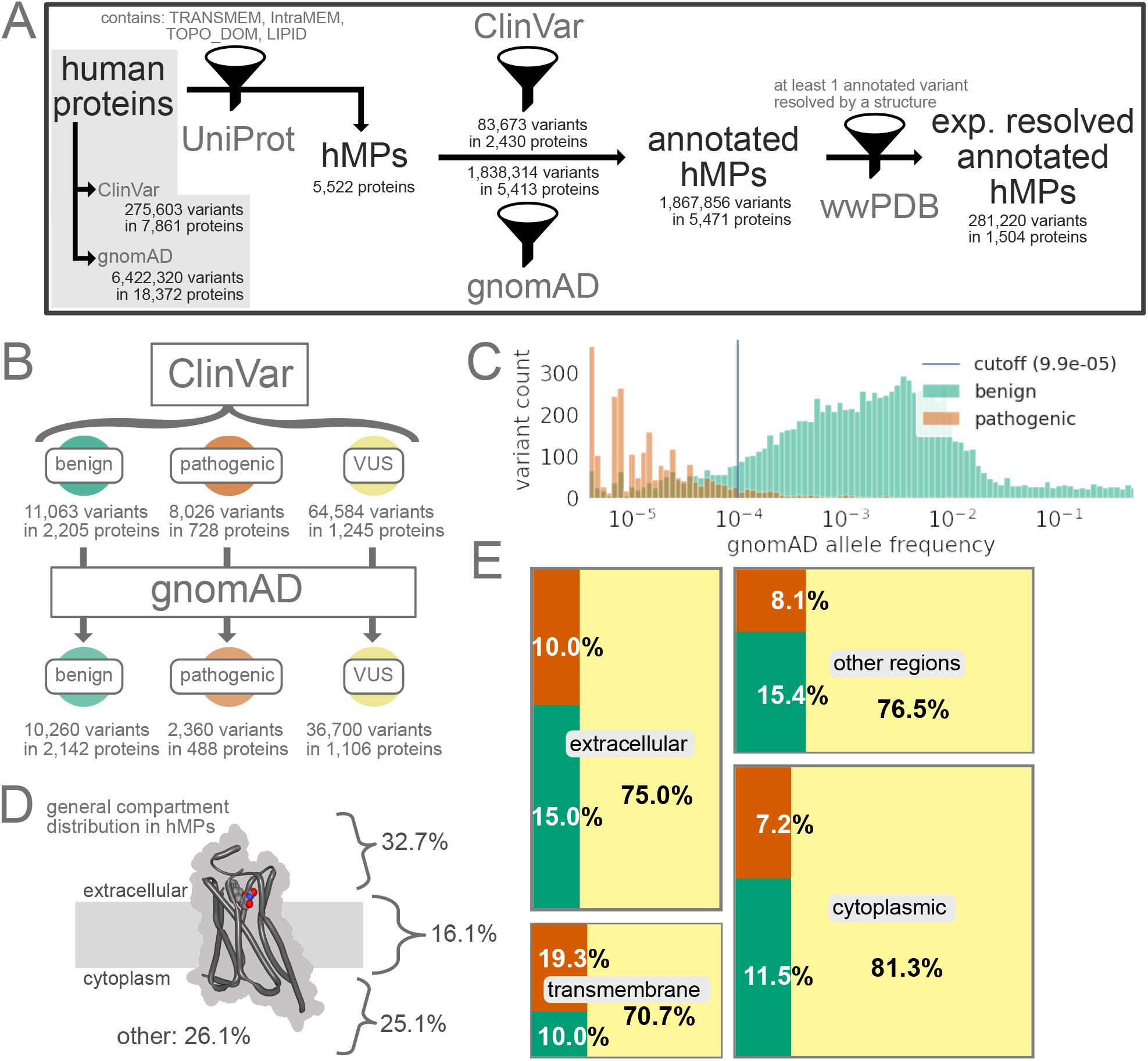
Overview of variants in human proteins and human membrane proteins (hMPs) in particular. A. Statistics on protein and variant annotations, ClinVar/gnomAD annotation, and availability of high-resolution structures mainly on human membrane proteins. B. ClinVar statistics for benign, pathogenic, and variants of unknown significance (VUS) and their coverage by gnomAD in human membrane proteins. C. gnomAD allele frequencies for all variants in human membrane proteins observed in gnomAD. Note that 71% of the pathogenic variants in ClinVar are not in gnomAD and hence missing from this analysis. D. Percentage of protein regions of different cellular elements across human membrane proteins. ‘Other’ includes compartments such as the lumen, mitochondrial matrix, and vesicular compartments. E. Distribution of ClinVar annotations (color scheme as in B) in different regions of the human membrane proteins.

### Filtering criteria for variant analysis of the 16 target proteins

Prior to analysis, we defined filtering criteria for the calculated ΔΔG and ΔΔE variants to obtain a set of variants with reliable scores. First, only variants for which both ΔΔG and ΔΔE calculations are available were selected. Second, *special* residues involved in disulfide bonds or known modified residues (such as those which bind covalent ligands or palmitoylated residues) were excluded. Further, variants with a low MSA sequence coverage of fewer than 50 sequences were excluded. Last, variants that have a positive wild-type Rosetta energy (E_res_) are excluded from further analyses as those residue conformers are likely to favor any substitution in order to reduce its energy, likely due to limitations of the Rosetta energy function. By applying all filters, we obtain a final set of 15 proteins with 220 pathogenic (*group A*) and 104 *group B* (benign and/or non-rare) variants, of which 42 are benign (see Table 3 and for the single filtering steps additional data at https://github.com/KULL-Centre/papers/tree/main/2022/hMP-Xray-Tiemann-et-al).

**Table 1:** Number of variants annotated by ClinVar (as benign, pathogenic, and VUS), gnomAD for all human proteins (including membrane proteins) and human membrane proteins exclusively. For membrane proteins, the total counts are further divided into each of the cellular regions they occur in. VUS (ClinVar) includes conflict variants.

|  | human proteins | human membrane proteins |  |  |  |  |
| --- | --- | --- | --- | --- | --- | --- |
|  | all | all | extracellular | cytoplasmic | transmembrane | other |
| total | 6,526,797 | 1,867,856 | 574,211 | 447,435 | 258,366 | 587,844 |
| benign | 36,770 | 11,063 | 3,489 | 3,544 | 961 | 3,069 |
| benign ∩ gnomAD | 33,944 | 10,260 | 3,214 | 3,326 | 908 | 2,812 |
| pathogenic | 21,107 | 8,026 | 2,327 | 2,233 | 1,863 | 1,603 |
| patho. ∩ gnomAD | 6,200 | 2,360 | 632 | 628 | 454 | 646 |
| VUS | 217,726 | 64,584 | 17,451 | 25,119 | 6,814 | 15,200 |
| VUS ∩ gnomAD | 116,325 | 36,700 | 9,907 | 14,693 | 3,574 | 8,526 |
| only gnomAD | 6,251,194 | 1,784,183 | 550,944 | 416,539 | 248,728 | 567,972 |

**Table 2:** Number of variants annotated by ClinVar or gnomAD, separated by their membrane protein category: single-pass, multi-pass, lipid-anchored, integral

| Membrane protein category<br>variant counts for | all |  | Single-pass |  | Multi-pass |  | Lipid-anchored |  | Integral |  |
| --- | --- | --- | --- | --- | --- | --- | --- | --- | --- | --- |
|  | all | transmembrane | all | transmembrane | all | transmembrane | all | transmembrane | all | transmembrane |
| total | 1,867,856 | 258,379 | 861,305 | 30,480 | 824,532 | 218,193 | 127,370 | 0 | 54,649 | 9,706 |
| benign | 11,063 | 961 | 4,960 | 138 | 4,794 | 744 | 817 | 0 | 492 | 79 |
| benign ∩ gnomAD | 10,260 | 908 | 4,634 | 132 | 4,455 | 699 | 690 | 0 | 481 | 77 |
| pathogenic | 8,026 | 1,863 | 2,328 | 77 | 4,741 | 1,719 | 544 | 0 | 413 | 67 |
| patho. ∩ gnomAD | 2,360 | 454 | 694 | 20 | 1,377 | 412 | 144 | 0 | 145 | 22 |
| VUS (ClinVar) | 64,584 | 6,814 | 25,707 | 693 | 29,677 | 5,784 | 5,311 | 0 | 3,889 | 337 |
| VUS ∩ gnomAD | 41,511 | 4,038 | 17,128 | 459 | 19,307 | 3,371 | 2,589 | 0 | 2,487 | 208 |
| only gnomAD | 1,784,183 | 248,741 | 828,310 | 29,572 | 785,320 | 209,946 | 120,698 | 0 | 49,855 | 9,223 |
| protein count | 5,796 |  | 2,330 |  | 2,762 |  | 561 |  | 143 |  |

**Table 3:** Overview of proteins analysed in depth, including Uniprot name, protein functional class, protein length and number of amino acids within the transmembrane (TM) region, number of group A (=pathogenic), group B (=benign and/or non-rare gnomAD) and benign variants before and after filtering, the sequence depth of the multiple sequence alignment (MSA) used by GEMME, the 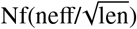 of the MSA, the AUC for our two predictors (ΔΔG and ΔΔE) and the MIM disease phenotype.

| protein info |  | length |  | before filtering |  |  | after filtering |  |  | GEMME |  | AUC |  | MIM |
| --- | --- | --- | --- | --- | --- | --- | --- | --- | --- | --- | --- | --- | --- | --- |
| name | class | all | TM | group A | group B | benign | group A | group B | benign | MSA depth | $Nf(neff/\sqrt{len})$ | $\Delta\Delta G$ | $\Delta\Delta E$ | phenotype |
| NPC1 | Transporter | 1278 | 277 | 60 | 36 | 13 | 44 | 12 | 5 | 1486 | 31.43 | 0.69 | 0.84 | Niemann-Pick disease |
| OPSD | GPCR | 348 | 161 | 67 | 10 | 2 | 41 | 6 | 2 | 1183 | 49.16 | 0.79 | 0.69 | Night blindness; Retinitis punctata albescens; Retinitis pigmentosa |
| GTR1 | Transporter | 492 | 261 | 56 | 9 | 4 | 42 | 7 | 2 | 1772 | 65.62 | 0.65 | 0.92 | Dystonia; GLUT1 deficiency syndrome; Epilepsy |
| AT2A2 | Transporter | 1042 | 204 | 12 | 9 | 7 | 8 | 5 | 4 | 1924 | 44.33 | 0.85 | 1 | Acrokeratosis verruciformis; Darier disease; |
| ACHB2 | Ion channel | 502 | 86 | 3 | 16 | 12 | 1 | 5 | 3 | 1789 | 57.23 | 0.8 | 1 | Epilepsy |
| CXB2 | Cell Junction | 226 | 83 | 60 | 23 | 8 | 33 | 16 | 5 | 1351 | 82.1 | 0.42 | 0.84 | Deafness; Bart-Pumphrey syndrome; Vohwinkel syndrome;... |
| S5A2 | Enzyme | 254 | 84 | 21 | 11 | 5 | 18 | 8 | 5 | 1177 | 60.18 | 0.63 | 0.9 | Pseudovaginal perineoscrotal hypospadias |
| MC4R | GPCR | 332 | 165 | 12 | 11 | 2 | 10 | 8 | 1 | 1265 | 54.02 | 0.66 | 0.9 | Obesity |
| AQP2 | Ion channel | 271 | 124 | 10 | 2 | 1 | 7 | 1 | 1 | 1326 | 62.36 | 1 | 0.86 | Diabetes insipidus |
| ACHA4 | Ion channel | 627 | 85 | 3 | 47 | 32 | 3 | 6 | 4 | 1107 | 27.84 | 0.33 | 0.94 | Epilepsy; Nicotine addiction |
| JAGN1 | Transporter | 183 | 84 | 5 | 4 | 3 | 4 | 3 | 3 | 196 | 13.45 | 0.42 | 0.92 | Neutropenia |
| SMO | GPCR | 787 | 147 | 1 | 25 | 6 | 1 | 7 | 1 | 823 | 17.7 | 1 | 0.86 | Curry-Jones syndrome; Pallister-Hall-like syndrome; Basal cell carcinoma |
| ABCG8 | Transporter | 673 | 126 | 1 | 35 | 11 | 0 | 15 | 3 | 86 | 2.14 | - | - | Gallbladder disease; Sitosterolemia |
| ABCG5 | Transporter | 651 | 127 | 1 | 27 | 9 | 0 | 0 | 0 | 31 | 0.81 | - | - | Sitosterolemia |
| GPT | Enzyme | 408 | 228 | 7 | 5 | 1 | 6 | 2 | 1 | 1812 | 64.45 | 0.5 | 0.92 | Congenital disorder of glycosylation; Myasthenic syndrome |
| FZD4 | GPCR | 537 | 206 | 5 | 13 | 6 | 2 | 3 | 2 | 1055 | 37.22 | 0.83 | 1 | Exudative vitreoretinopathy; Retinopathy of prematurity |
| total |  |  |  | 324 | 283 | 122 | 220 | 104 | 42 |  |  | 0.64 | 0.82 |  |

### Definition of ΔΔG and ΔΔE thresholds and quadrant classification

To analyze the variants in terms of their ΔΔE and ΔΔG scores, we defined cutoffs for each method based on the optimal ROC value (tradeoff of high specificity versus high sensitivity) to separate *group A* (pathogenic) and *group B* (benign and/or non-rare) variants, in a similar fashion as done for gnomAD allele frequency above. Next, we defined four categories dependent on their position along the ΔΔE and ΔΔG axes:

I. (I) *Quadrant (I)* variants have high ΔΔE and ΔΔG and are likely to cause loss of function via loss of stability;
II. (II) *Quadrant (II)* variants have high ΔΔE and low ΔΔG and cause loss of function for other reasons than loss of stability;
III. (III) *Quadrant (III)* variants have low ΔΔE and low ΔΔG and are from a structural and evolutionary perspective expected to be tolerated;
IV. (IV) *Quadrant (IV)* variants have low ΔΔE and high ΔΔG and are from an evolutionary perspective expected to be tolerated but from a structural perspective expected to cause instability;

### Definition of protein regions

For analysis purposes, we assigned residues into different regions, based on their solvent accessibility, and their positioning within the membrane (TM region). Relative solvent-accessibility was calculated using DSSP (51, 52) with a cutoff of 0.3, placing residues with a smaller value into the category of *buried* residues. The positioning within the membrane was obtained as described above. We can thereby divide the protein into four regions:

- *Buried*: residues with a DSSP < 0.3 and that are placed outside the membrane; this cutoff places residues within contacts as buried although they might be close to the surface of the protein
- *Solvent-accessible:* residues with a DSSP ≥ 0.3 and that are placed outside of the membrane
- *TM-region* ⋂ *buried*: buried residues that are placed within the membrane
- *TM-region* ⋂ *solvent-accessible*: solvent-accessible residues within the membrane

For additional analysis, we divide residues by whether they are oriented toward the lipids or not. To assess whether a residue faces toward the lipids, we used a dedicated Rosetta function that returns a true or false value (50). Most of those residues are within the transmembrane region but there can be exceptions that non-transmembrane residues (either solvent accessible or buried) can face the lipids by “dipping” into the membrane plane. For Fig. S4C and D, we expended the regions above by 3 more which are subsections of the above (with overlaps, see Fig. S4):

- *TM-region* ⋂ *lipid-facing* ⋂ *buried*: buried residues within the membrane that are oriented towards the lipids
- *TM-region* ⋂ *lipid-facing* ⋂ *solvent-accessible*: solvent-accessible residues within the membrane that are oriented towards the lipids
- *Others*: combination of residues that are rare and few in number, such as *TM-region* ⋂ *solvent-accessible, lipid-facing* ⋂ *solvent-accessible* or *lipid-facing buried*

### Utilized software

- python3, incl. following third party libraries

– adjustText
– Biopython
– circlify
– matplotlib
– numpy
– pandas
– seaborn
– scipy
– sklearn
– squarify
– xmltodict
– nglview
- Rosetta version 2021.31+HEAD.c7009b3115c (c7009b3115c22daa9efe2805d9d1ebba08426a54, default.linuxgccrelease mode)

– mp_span_from_pdb (54)
– rosetta_scripts (60)
Tiemann, Zschach, Lindorff-Larsen and Stein
– FastRelax (61, 62)
– MembraneMover (54)
– cartesian_ddg (12, 14)
– energy function: franklin2019 (35) + cart_bonded=0.5 + fa_water_to_bilayer=1.5
- Scripts available at https://github.com/KULL-Centre/papers/tree/main/2022/hMP-Xray-Tiemann-et-al

– hMP statistics (hMP_stats.ipynb)
– ΔΔG pipeline benchmark (MP_ddG_benchmark.ipynb)
– Xray subset calculations (Xray_subset-calc.ipynb)
– Xray subset analysis (Xray_subset-ana.ipynb)
- Pipelines

– PRISM_tools (https://github.com/KULL-Centre/PRISM/software/rosetta_ddG_pipeline, release version v0.1.1)
– PrismData, FillVariants and struc_select_sifts (https://github.com/KULL-Centre/PRISM/software/scripts, release version v0.2.2)
- Others

– overleaf.com
– Affinity Designer

### Limitations of this study

We note several limitations that should be considered when interpreting the results. Our general observations and conclusions on membrane proteins and their classes are limited by the available data and proteins, partly due to our choice to only analyze experimental structures with annotated pathogenic and benign variants.

Several membrane proteins, for example channels and cell junction proteins, function as (homo-) oligomers. In this study, we used structures of the individual proteins for our stability calculations and thereby may miss destabilizing variants in interfaces. Those variants are more difficult to interpret using stability calculations due to the lack of contacts that are affected by stability. In addition to missing interactions, conformational changes of the structure or different conformations might alter ΔΔG values, and several stabilizing variants can be explained due to missing interaction partners in these structures (e.g. R135W, R135L, and G121V in OPSD are missing either the ligand or an intracellular binding partner).

In general, we do not expect variants to lead to complete protein unfolding but rather a partial unfolding, which allows recognition by the protein quality control system. Due to limited available experimental data, we are not able to differentiate stages of unfolding, which might affect the accuracy of the ΔΔG calculations. Furthermore, our membrane protein dataset is mainly alpha-helical, which is also true for most human membrane proteins; however, the stability score function was parameterized and benchmarked on bacterial proteins, which are often beta barrels and might fold differently compared to their helical counterparts. When we compared experimental and computational ΔΔG values, we obtained a Spearman rank correlation coefficient of 0.46, leaving uncertainty about the predictability of the extent of loss of stability. It is worth noting that the correlation between two sets of experimental ΔΔG measurements in GlpG (57, 58) shows a Spearman correlation of 0.65, and when correlating all experimental datasets with at least 12 overlapping variants we obtain a mean Spearman correlation of 0.6. Preferences for specific amino acid properties in certain environments such as the membrane might be biased by their values within the respective scoring function. Our results also depend on how different protein regions are defined.

Co-evolutionary sequence conservation measurements cannot give direct insights into the mechanism that causes pathogenicity. Variants labeled *lose function* here may instead exhibit the more rare gain of function instead. Furthermore, our calculations of ΔΔE scores depend on the MSA, and we note that using a different MSA, e.g., by changing the E-value cutoff, could shift some of the ΔΔE values from tolerated to not tolerated.

The filters we apply on the variants are chosen based on literature and experience. A more detailed analysis on the effects of this filtering (and their cutoffs) with a larger dataset of variants is needed. For now, we applied AUC calculations on each of the filtering steps to address a potential bias (see Supplementary Material).

Finally and as already discussed above, we combine benign and/or non-rare gnomAD variants into group B. This should be taken into account when interpreting the results and especially when investigating outliers of group B, as those could be variants of unknown significance.

## RESULTS AND DISCUSSION

### Variant annotations in human membrane proteins

We first set out to obtain an overview of the presence and properties of missense variants in human membrane proteins. We searched Uniprot (42) for keywords like TRANSMEM (see Methods and Materials for details) and used the results to define a list of 5,522 proteins that are thought to be embedded in the membrane (Fig. 1A). We subsequently searched the gnomAD (44) and ClinVar (43) databases for missense variants in the genes encoding these proteins (see Methods and Materials for additional details). gnomAD is a database aggregating the variants observed in ca. 150,000 exome and genome sequences, and thus provides a relatively unbiased view of the variants that are present in the human population (44). ClinVar is a database containing, among other things, missense variants that have been categorized as benign, pathogenic, or *variants of uncertain significance* (VUS), the latter indicating that the pathophysiological consequences of the variant are not clear (43). We obtain almost 1.9 million variants in total for human membrane proteins, which makes up 29% of all human protein variant annotations (see Table 1). Almost all (98.1%) membrane proteins have at least one variant in gnomAD, and about half (44.0%) have at least one variant in ClinVar (Fig. 1A). Across the two datasets, we find 1.9 · 10^6^ unique variants in 5,471 membrane proteins. We excluded synonymous, silent, or deletion variants, which make up 0.3% (5,403 variants) from any further analysis. Nearly all (99.1%) of the non-synonymous variants are either from gnomAD or are assigned as variants of uncertain significance in ClinVar, and only 19,089 of the 1.9 million variants have an assigned status of being pathogenic or benign (Fig. 1A and Table 1), highlighting the scope of the problem of determining variant effects. 38% of all human pathogenic variants are found within membrane proteins (see Table 1), underlining the importance of method development suited for this protein class.

Variants that are pathogenic are expected to be depleted in the human population compared to those that are benign, and indeed we find a clear separation of the distributions of allele frequencies between the two classes (Fig. 1C). We also observe that while 92.7% of benign ClinVar variants have been observed in gnomAD, this is only true for 29.4% of pathogenic variants (Fig. 1B, Table 1). The separation in the distribution of allele frequencies between pathogenic and benign variants suggests that variants with allele frequencies > 9.9 · 10^-5^ are more likely to be benign than pathogenic (Fig. 1C, cut-off calculated from the receiver-operator characteristic (ROC) analysis, see Methods and Materials). While the allele frequency in gnomAD appears to be a good predictor of pathogenicity (area under the curve (AUC) of 0.96; Fig. S1, with similar results for all human proteins; AUC=0.95), we note that this result should be taken with some caution. First, many ClinVar variants are not found in gnomAD (Fig. 1B), limiting the practical utility. Second, since the presence in gnomAD might have been used to assign (lack of) pathogenicity, it is difficult to ensure that the two sets of data are independent.

We analyzed in which regions of the membrane protein structures the ClinVar (Fig. 1D,E) and gnomAD (Table 1) variants are located. We find that most variants are found in soluble domains, though this is likely due to the fact that these regions makes up 83% of membrane proteins (Fig. 1D,E). Notably, though, we find that while the numbers of known benign and pathogenic variants are similar in the different types of soluble regions, there appears to be an almost two-fold excess of pathogenic variants compared to benign variants in the transmembrane regions (Fig. 1E, Table 1). While we cannot exclude that this enrichment is in part due to an increased focus on the transmembrane region in clinical research, we suggest—in line with previous work (63, 64)—that this observation also reflects a decreased mutational tolerance of the transmembrane region.

Membrane proteins are typically defined by their interaction and/or location within the membrane. As not all of them are located to a similar degree inside the membrane, we divided the complete dataset of 5,796 membrane proteins into their categories as being *single-pass, multi-pass, lipid-anchored*, or *integral* membrane protein (see table 2). We find that most proteins are single-(40.2%) or multi-pass membrane proteins (47.7%) and also most of the variants are found in these categories (46.1% and 44.1% of 1,867,856 variants). Looking into the transmembrane region, we see the previously described enrichment of pathogenic variants especially for multi-pass membrane proteins. This makes the multi-pass membrane protein category especially interesting for further studying of the role of residues within this region.

To gain a better understanding of the mechanisms causing benign or pathogenic variant consequences, we mapped the variants onto known protein structures. Despite recent advances in protein structure prediction (65) and analysis using computational methods (66, 67), we decided to focus our work on experimentally determined structures. Specifically, we searched the protein databank (68) for structures with at least one variant in the resolved part of the protein structure and found that 27.5% of all annotated human membrane proteins have at least some part resolved and that 15.1% of the total set of variants are found in the region covered by these structures (Fig. 1A, additional data at https://github.com/KULL-Centre/papers/tree/main/2022/hMP-Xray-Tiemann-et-al). Of the 281,220 variants found in resolved regions, only 2.2% of those (6,119 variants) have been assigned as benign (2,089 variants, 18.9% of total benign variants in membrane proteins) or pathogenic (4,030 variants, 50.2% of total pathogenic variants in membrane proteins) (Table S1).

### Computational assessment of stability and evolution shows loss of function due to loss of stability for ca. 62% of disease variants in selected proteins

To examine the importance of changes in protein stability in membrane proteins for causing loss of function and disease, we analysed a smaller set of proteins in more detail. Specifically, we searched for proteins that had at least one pathogenic and one benign variant in the transmembrane region. As our aim was to use the Rosetta software to predict changes in thermodynamic stability, we focused on protein structures that had been determined via X-ray crystallography as Rosetta has been developed and benchmarked most extensively on such structures. These requirements narrow down the set to 16 proteins: six transporters, three ion channels, four GPCRs, two enzymes and one cell junction protein (Table 3 and additional data at https://github.com/KULL-Centre/papers/tree/main/2022/hMP-Xray-Tiemann-et-al). These 16 proteins represent different types of membrane proteins with diverse functions, structures, and involvement in different diseases. Of note, these proteins belong to the class of multi-pass transmembrane proteins and the secondary structure of these proteins is mostly α-helical (> 50%), while unstructured regions or extended strands add up to 27% (Fig. S2).

Inspired by previous analyses of soluble proteins, we investigated these membrane proteins in terms of structural stability and sequence conservation. Specifically, we developed and benchmarked a revised Rosetta protocol for stability calculations of membrane proteins (see Methods and Materials and Supplementary Material). We used this method to calculate the change in thermodynamic stability (ΔΔG) upon single amino acid substitutions. In each case, we selected a high-resolution structure (Table 3), and removed any co-crystallised molecules. We also constructed multiple sequence alignments (MSA) of each protein and used GEMME (Global Epistatic Model for predicting Mutational Effects) (36) to estimate the evolutionary effects of the variants. Specifically, we calculated a normalized score (ΔΔE) with ΔΔE ≈ 0 corresponding to substitutions that—in light of evolution—appear well tolerated, and ΔΔE ≈ 1 for variants that—based on the evolutionary record—are rare or absent, and expected to cause loss of function. In analyses of soluble proteins, we have previously found that a high value of ΔΔE is a good predictor for a variant to cause loss of function and that variants with both high ΔΔE and ΔΔG are likely to cause loss of function via loss stability and cellular abundance (18).

We calculated ΔΔG and ΔΔE for all variants that have been observed in humans and where the wild-type residue was resolved using X-ray crystallography. Further, we did not analyze variants at positions where the Rosetta energy function suggested a potential incompatibility between the experimental structure and the Rosetta energy function (e.g. disulfide bridges (filter II) or residues with a positive energy where mutations are likely more tolerated by default (=E_res_, filter IV).), and variants at positions with 499731 50 sequences in the multiple sequence alignment (see Methods and Materials, Table 3 and Supporting Data at https://github.com/KULL-Centre/papers/tree/main/2022/hMP-Xray-Tiemann-et-al, table *2022_11_11-count_hMP_anno_nonsyndel_PDB_publish*, tab *X-ray_set_app* for the variant loss at each of the sequential filtering steps and further information). After this quality control, we retain 220/324 pathogenic and 42/122 benign variants and lose one protein (ABCG8) as it does not have any variants left after filtering. We thus analyzed two sets of variants: A is the set of 220 variants that are described as pathogenic in ClinVar and B is the set of 104 variants that are either assigned as benign in ClinVar and/or non-rare gnomAD variants that, as discussed above, are more likely to be benign than pathogenic as their allele frequencies in gnomAD are > 9.9 · 10^-5^ (Fig. 1 and Fig. S1). In what follows we refer to *group B* as benign, but note that among the 104 variants in *group B* only 42 are classified as benign in ClinVar and the remainder comes from gnomAD. To get an indication of the influence of this filtering process, we performed all AUC measurements also on the respective filtering steps/subsets (see Extended supplemental figure collection 1 worksheet tab *X-ray_set_app_AUC*).

To quantify how well ΔΔE and ΔΔG distinguish between the two classes of variants, *group A* (pathogenic) and *group B* (benign and/or non-rare), we constructed a ROC curve and calculated the AUC as a measure of how well each of the two scores can predict pathogenicity (Fig. 2A). Of note, to ensure our results are not biased by the limited dataset, we performed a leave-one-protein-out calculations when performing the ROC curves and their derived cutoffs, giving us mean values with standard deviation for the AUC and mean, min and max cutoffs values. In the following, we report for AUC by AUC_l1po_ = mean AUC ± std. Variant counts in the quadrant (and their respective percentages) are determined from the total cutoffs and the leave-one-protein-out calculations. In the latter, variants that are located inside the min to max leave-one-protein-out cutoff values are considered as “gray” and contribute to the standard deviation of the reported percentages. Looking at out complete data, we find that both ΔΔE and ΔΔG can separate the *group A* and *group B* variants, though ΔΔE, as expected, performs better than ΔΔG (AUC 0.82 vs. 0.64; ΔΔE AUC_l1po_ = 0.82 ± 0.01, ΔΔG AUC_l1po_ = 0.62 ± 0.01). This is in line with previous observations for soluble proteins (6, 7, 9, 11) and the hypothesis that many, but not all, pathogenic variants are destabilized so that ΔΔG calculations can capture these pathogenic variants, but not those caused by other mechanisms for loss/gain of function.

**Figure 2:**
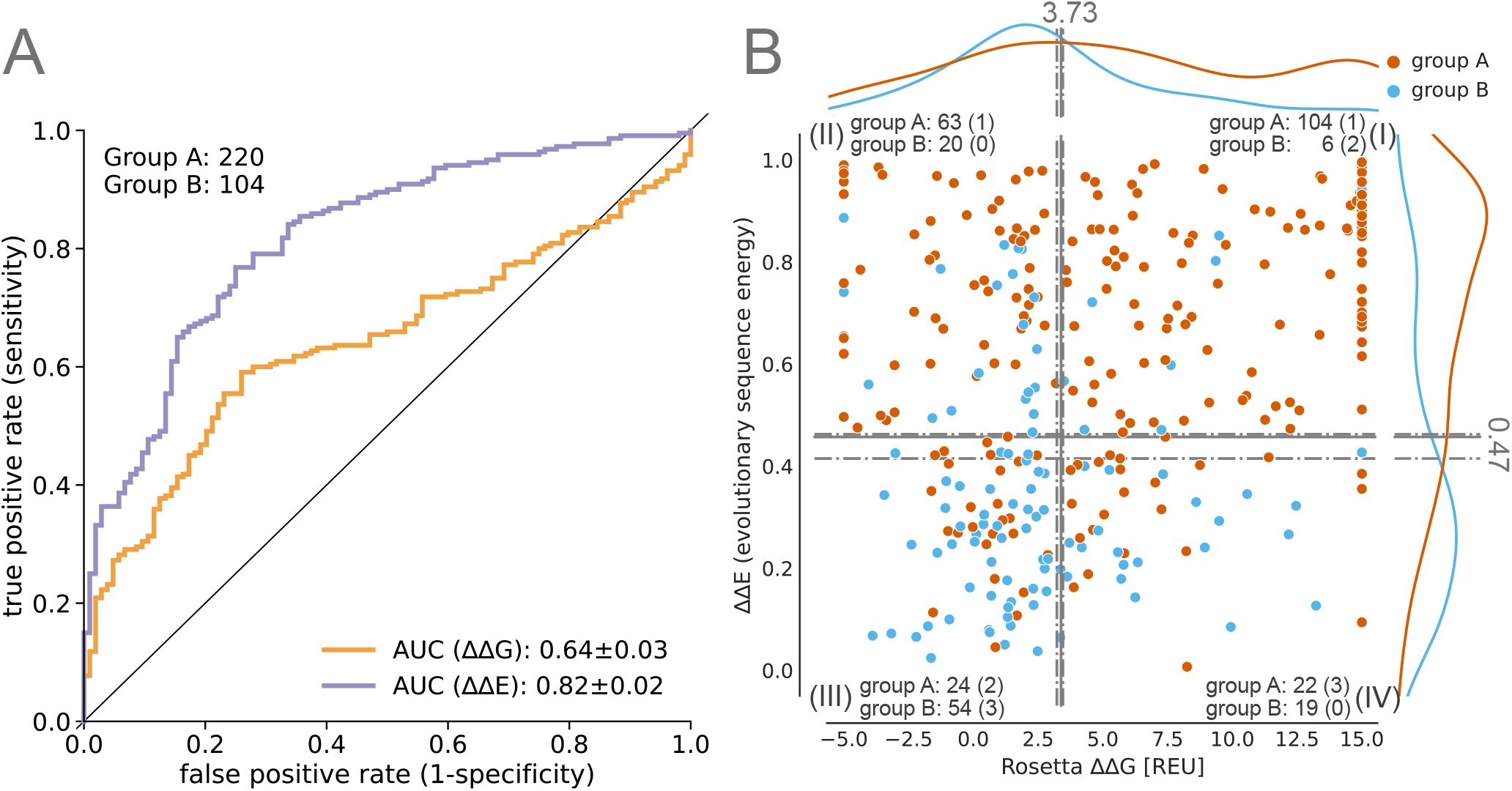
Correlation of ClinVar variants with computational predictors for human membrane proteins. (A) ROC curves of Rosetta ΔΔG and GEMME ΔΔE with variant counts for each group. AUC standard errors were determined by bootstrapping. (B) Distribution of *group A* (pathogenic) and *group B* (benign and/or non-rare) along ΔΔG and ΔΔE landscape and separation into quadrants by mean, max and min optimal cutoff obtained from leave-one-protein-out ROC curves. The counts of each group are shown for the quadrants, the respective number of variants in the uncertain area in brackets.

We further analyzed the *group A* (pathogenic) and *group B* (benign and/or non-rare) variants in terms of their ΔΔE and ΔΔG scores (Fig. 2B). To simplify the discussion, we analyze the variants in terms of whether ΔΔE and ΔΔG are low or high, with respect to cutoffs from the ROC analysis (see Methods and Materials section). This analysis separates the variants into four quadrants with only a few variants (14.2%, mean_l1po_ = 13.6 ± 5.9%) falling in the quadrant of low ΔΔE and high ΔΔG (Fig. 2B (IV)), which is comparable to previous observations for soluble proteins (18, 19). The three remaining quadrants correspond roughly to:

1. (I) Variants that cause loss of function via loss of stability (high ΔΔE and high ΔΔG),
2. (II) Variants that cause loss of function for other reasons than loss of stability, such as substitutions at key functional sites (high ΔΔE and low ΔΔG),
3. (III) Variants that are expected to be tolerated both from a structural and evolutionary sequence perspective (low ΔΔE and low ΔΔG).

As expected, we find that most *group B* (benign and/or non-rare) variants have low ΔΔE (75%, mean_l1po_ = 73.1 ± 2.9%) and 73% (mean_l1po_ = 75.0 ± 3.9%) of those also low ΔΔG, whereas most pathogenic variants *(group A)* have large ΔΔE (76.8%, mean_l1po_ = 76.8 ± 0.9%, see Fig. 2B). Among the 167 (2 in gray area) pathogenic variants that have high ΔΔE values, we find that 62.1% (mean_l1po_ = 61.5 ± 0.6%) also have high ΔΔG values, suggesting that loss of stability plays an important role for disease in the 15 investigated membrane proteins.

We observe a number of pathogenic variants with very negative ΔΔG that indicates a stabilizing effect on the structure. As also recently shown by (69), gain of function variants can lead to pathogenicity, those variants we observe might be explained in a similar way. To confirm, more information and benchmark is needed.

### Pathogenic variants in GPCRs, especially in the transmembrane region, lose function mostly by loss of stability, while this is less prominent in transporters or other protein classes

Our data set contains several members of the main membrane protein classes, namely five transporters (98 *group A*/pathogenic + 62 *group B* (benign and/or non-rare variants)), three ion channels (11 + 15 variants), four GPCRs (54 + 28 variants) and two enzymes (24 +13 variants) (Table 3). We examined the results from our computational predictors to probe for class-specific trends. In all four classes, evolutionary conservation predictions (ΔΔE) have a high AUC (>0.8), similar to the analysis with all proteins combined (Table S2). Focusing on the transmembrane region, we find a very high AUC of 0.97 (AUC_l1po_ = 0.98 ± 0.02) for variant classification of transporters (underlying datapoints: 44 pathogenic *group A* + 10 *group B* variants). Interestingly, we find that for GPCRs loss of stability is the main cause of pathogenicity, as indicated by an increased AUC (0.79, AUC_l1po_ = 0.77 ± 0.05) for ΔΔG predictions (Fig. 3A and Table S2) compared to the complete data set with 15 proteins (AUC=0.64, AUC_l1po_ = 0.62 ± 0.01, Fig. 2). This is even more prominent for variants located within the transmembrane region (AUC=0.81, AUCl1po = 0.81 ± 0.03).

**Figure 3:**
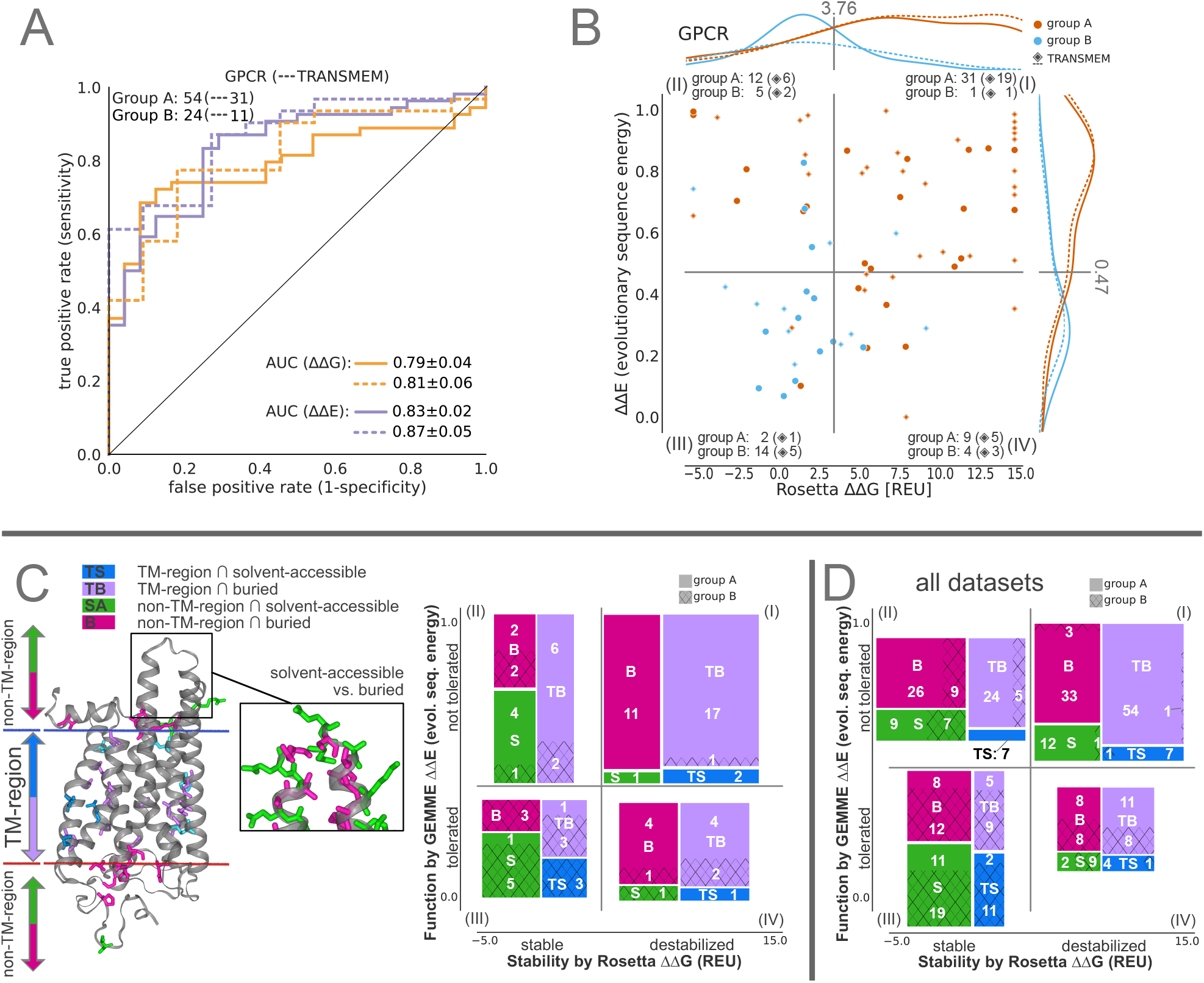
Analysis of GPCR variant classification performance and structural differences of variant effects. (A) ROC curve and (B) distribution of ΔΔG versus ΔΔE plotted for GPCRs for all variants and only for variants within the transmembrane region (dashed line and diamond symbol). (C) [left] Illustration of the different residue categories used in this work on OPSD, namely, whether they are inside (blue, and violet) or outside of the transmembrane (TM) region (green and pink), and whether they are solvent-accessible violet and pink or buried blue and green. The structure on the left shows disease-associated variants, while the excerpt on the right illustrate solvent-accessible vs. buried, more generally and are for illustration purposes not restricted to a disease relationship. For more details on how the classes were assigned, see Methods and Materials. On the right, the variant counts in the four quadrants, separated by their position in the proteins, are shown for *group A* (pathogenic, full) and *group B* (benign and/or non-rare, hashed) variants. (D) Variant counts as seen in (C) are shown summed over all 15 proteins.

In the transmembrane region of GPCRs, 77.4% (mean_l1po_ = 74.2 ± 25.8%) of the pathogenic variants have high ΔΔG values (Fig. 3B), suggesting that their pathogenicity is due to loss of stability. When separating the proteins into specific regions, namely by whether they are buried, solvent-accessible, and are within and outside the transmembrane regions (Fig. 3C), we see that those pathogenic variants that lose function via loss of stability are typically buried (Fig. 3C). In contrast, solvent-accessible pathogenic variants are not found to lose function due to loss of stability, and variants located in those regions are more likely to be tolerated (11.1% of *group A*/pathogenic compared to 44.4% of *group B* variants). Within the transmembrane region, most variants (90% *group A* and 97% for *group B*/benign and/or non-rare) are buried, in contact with other residues. Looking at pathogenic variants that lose function due to other reasons than loss of stability (quadrant (II)), variants in GPCRs are more often within the transmembrane (Fig. 3C) compared to all datasets (Fig. 3D), where we see a larger proportion of variants in buried sites in extracellular or intracellular environments. When we further divide residues into whether they are facing the lipid bilayer or not, we see that most of those pathogenic variants within the transmembrane region are facing the lipids while being in contact with other residues as indicated by their buried-ness (supp. Fig. S4) in contrast to their likely benign counterpart that is seen to be more solvent accessible.

Next, we focused on individual proteins and examined the location and potential mechanism behind disease variants in one GPCR and one transporter protein. We used the calculated values of ΔΔE and ΔΔG to aid in a structural analysis of the disease variants in rhodopsin (OPSD; Fig. 4A–4C) and a glucose transporter (GTR1; Fig. 4D–4F). We examined the structures of the two proteins to find the residues that interact with ligands or co-factors and searched the literature to find residues that are known to be key to function. We find that many disease variants are located at these residues, suggesting that they directly disrupt function, and some of them also decrease stability. For example, in OPSD we find a number of disease mutants at residues that interact with the retinal co-factor as well as residues in e.g. the so-called ionic lock (71) (Fig. 4A and B). Similarly, many disease variants in GTR1 are located at sites known to interact with a chloride ion that is important for function (70), the sugar molecule, known inhibitors (72), or residues known to affect transport (70) (Fig. 4C and D).

**Figure 4:**
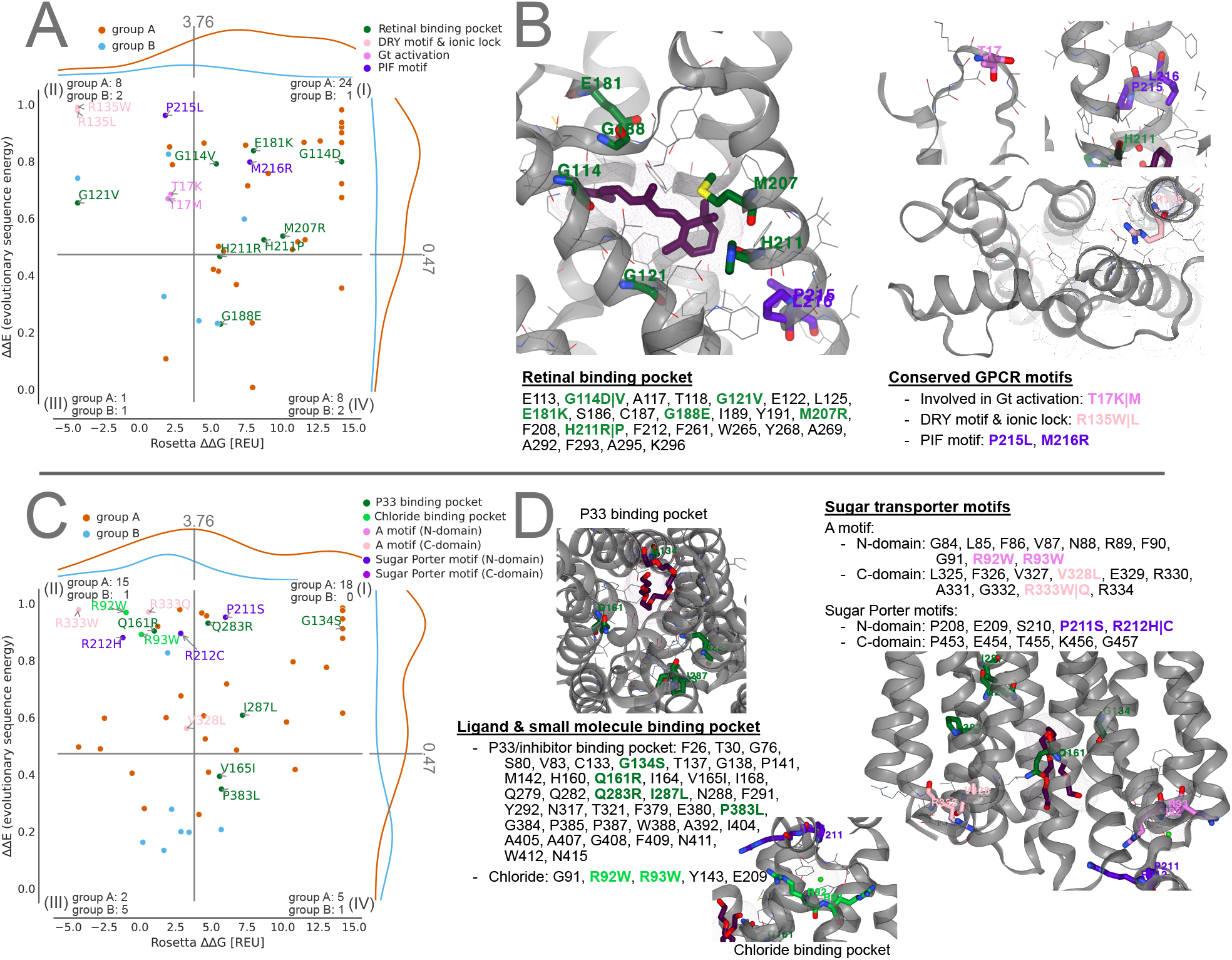
Mapping pathogenic variants to known motifs and binding pockets. A-C show OPSD variants which lie within the retinal binding pocket and conserved GPCR motifs. D-F show GTR1 variants within proximity of a PEG molecule (P33), matching previously identified inhibitor positions (70), and the chloride binding side, along with conserved sugar transporter motifs.

Looking across the two proteins (Fig. 4), most of the high-ΔΔE, low-ΔΔG disease variants are found at residues that have known functional roles. We expect such variants in quadrant (II) to lose function due to other reasons than stability (18). This also includes variation in residues in close proximity to ligands and interaction partners, which were not included in our stability calculations. Further, we find that many of the disease variants that are not located at known functional sites have both high values of ΔΔE and ΔΔG, suggesting that these variants instead disrupt the stability of the folded state.

### Correlating physicochemical changes with variant effects

We examined the dataset containing all 15 proteins and the amino acid properties within the four quadrants, where quadrant (I) contains destabilized and quadrant (II) stable variants while both quadrants (I) (II) are—in light of evolution—not tolerated. Quadrants (III) and (IV) are evolutionary tolerated, but quadrant (III) contains stable and quadrant (IV) destabilized variants (see Fig. 5A and B, and Methods and Materials for a more detailed quadrant definition). Across all quadrants, hydrophobic amino acids are most commonly observed (wild-type, 33%; target, 37%), which can be explained by the general preference for hydrophobic residues in membrane proteins, especially within the TM region (23). Almost 65% of the *group B* (benign and/or non-rare) variants located in quadrant (III) have, as expected, the same amino acid property for wild-type and target (35.1% remain hydrophobic, 21.1% charged, 8.8% polar). For the pathogenic variants that lose function due to loss of stability (quadrant (I)), we see greater changes in physicochemical properties among those substitutions (Fig. 5A). Interestingly, in quadrant (II), where variants lose function due to other reasons than stability, we see mainly hydrophobic target amino acid types (54.8%, with one third coming from charged to hydrophobic substitutions).

**Figure 5:**
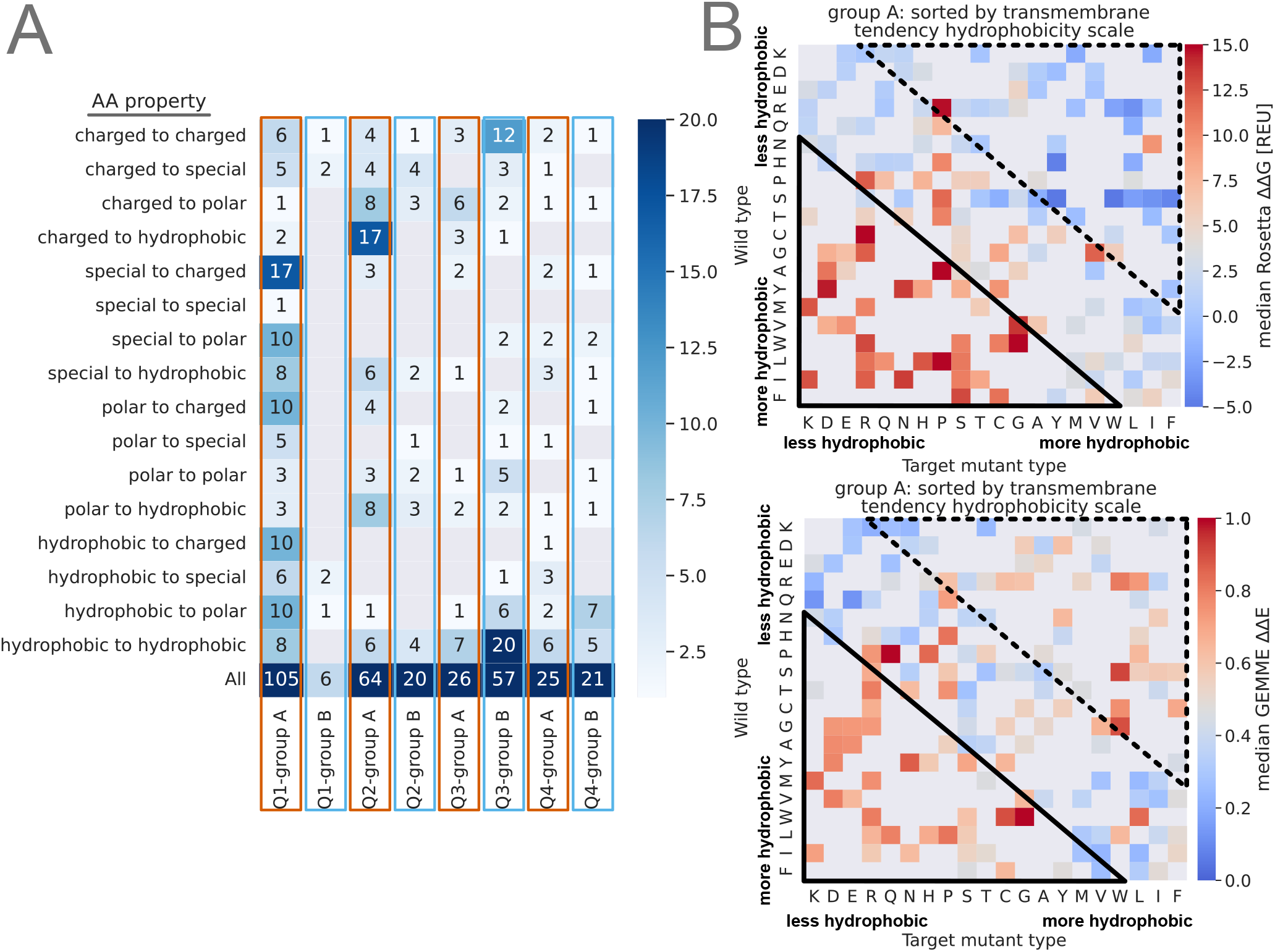
Variant effects depend on the chemical nature of the wild-type and target amino acids. (A) Substitution amino acid property counts for wild type and variant per quadrants and groups. Quadrant (I) contains destabilized and evolutionary not tolerated variants; quadrant (II) stable and not tolerated; quadrant (III) stable and tolerated and quadrant (IV) destabilized and evolutionary tolerated variants (Methods and Materials). (B) Heatmap of target versus wild-type amino acid, colored by their median values ΔΔG (top) and ΔΔE (bottom) for pathogenic variants sorted by the hydrophobicity ‘transmembrane tendency’ scale by (73). The rectangles show the enrichment of variants with high (solid line) and low (dashed line) ΔΔG or ΔΔE. Median values of ΔΔG (destabilized) and ΔΔE (not tolerated) are red, while low values are blue; missing data are colored gray.

Inspired by the enrichment of pathogenic variants in the transmembrane region (Fig. 1E) that is enriched with hydrophobic residues (23), we analyze substitutions by physicochemical properties. Specifically, we calculated the median score for ΔΔG and ΔΔE for each combination of wild-type and target amino acids and arranged the amino acids by hydrophobicity (73) (Fig. 5B). For variants where the target residue is more hydrophobic (e.g. Arg to Leu, Arg to Trp, or Asp to Tyr variants), we indeed see a different pattern when looking at the median stability values (ΔΔG) compared to the median ΔΔE values. These variants appear to be tolerated by protein stability, but not by evolution (Fig. 5B, dashed upper rectangle). In contrast, variants changing the residue to be less hydrophobic are indicated as not tolerated by evolution and destabilising (Fig. 5B, solid lower rectangle).

## CONCLUSIONS

Here, we present an analysis of missense variants and their properties within human membrane proteins. We identified 1.9 · 10^6^ unique variants in 5,471 proteins of which 99.1% are of uncertain significance, and only 19,089 have been classified as pathogenic or benign. Additionally, we see an almost 2-fold excess of pathogenic variants compared to benign variants in the transmembrane regions, which make up only 16.1% of the proteins.

We have examined the importance of changes in membrane protein stability for causing loss of function. We analysed 15 proteins and calculated the change in thermodynamic stability (ΔΔG) and evolutionary conservation (ΔΔE). Our ROC analysis shows good performance in separating benign from pathogenic variants by their sequence conservation (AUC 0.82 for ΔΔE), and we find that for our 15 analyzed transmembrane proteins ca. 62% of the pathogenic variants cause loss of function via loss of stability. This indicates that loss of stability indeed plays an important role for disease variants in membrane proteins, in line with previous findings on soluble proteins, although this needs to be confirmed with studies on larger datasets. In the 15 selected proteins, we observe that most variants have a hydrophobic wild type (33%) or target (37%) amino acid type and that almost 65% of benign and/or non-rare variants that are likely tolerated as assessed by both ΔΔE and ΔΔG do not change their amino acid type. Among pathogenic variants that lose function due to loss of stability, substitutions to charged, polar or hydrophobic are more prominent, while we observe substitutions from more hydrophobic to less hydrophobic residues in variants that lose function due to other reasons than stability.

When analyzing the different classes of membrane proteins, we observe for transporter proteins that pathogenic variants in the transmembrane region have an AUC of 0.97 for ΔΔE, and loss of stability does not appear to be the predominant factor in loss of function for the cell junction protein we examined. In contrast, pathogenic variants in GPCRs lose function mainly via loss of stability (AUC of 0.79 for ΔΔG). We therefore suggest that pathogenic variants lose function via loss of stability more often in the transmembrane region of GPCRs than in the other protein classes we examined.

From a more detailed inspection of individual proteins, we found that most of the high-ΔΔE, low-ΔΔG disease variants are located at positions that have known functional roles, while many of the disease variants that are not located at functional sites have both high values of ΔΔE and ΔΔG, suggesting that these variants instead disrupt the stability of the folded state.

Our observations underline the importance of stability and the loss thereof in disease-causing variants of membrane proteins and thereby show how computational tools can aid in interpreting molecular mechanisms that underlie disease. Such functional understanding may help address the substantial challenge of classifying VUS (74). Given the limited number of variants and proteins within this study, utilizing recent advantages like the large excess of experimental structures derived from electron microscopy or computational models from e.g. AlphaFold (75) could enable a broader analysis. We include the collection of population and ClinVar variants for the entire human proteome to facilitate such studies on membrane proteins and beyond.

## AUTHOR CONTRIBUTIONS

KL-L and AS conceived the original idea and supervised the project. JKST retrieved and processed all data with contribution by HZ who extracted and processed the ClinVar and gnomAD data. JKST and AS designed the Rosetta pipeline framework and JKST implemented and benchmarked it with KL-L and AS. JKST performed all calculations and processing of the data. JKST analyzed the data and interpreted the results with KL-L and AS. JKST, KL-L and AS wrote the manuscript with input from HZ.

## ACKNOWLEDGMENTS

We thank Matteo Cagiada for providing an automatic pipeline for GEMME calculations and Kristoffer Enøe Johansson for resourcing us with his alignment and merging implementations. Additional thanks go to Julia Koehler Leman for helpful discussion regarding membrane protein implementations in Rosetta. This study was funded by the Protein Interactions and Stability in Medicine and Genomics (PRISM) centre funded by the Novo Nordisk Foundation (NNF18OC0033950, to AS and KL-L) and a grant from the Lundbeck Foundation (R272-2017-4528, to AS). We acknowledge access to resources from the Department of Biology’s core facility for biocomputing.

## DATA AVAILABILITY

All scripts are available at https://github.com/KULL-Centre/papers/tree/main/2022/hMP-Xray-Tiemann-et-al and as stated in the ‘Utilized software’ section.

All parsed data fromUniprot, ClinVar, and gnomAD for the human proteome are available at https://sid.erda.dk/sharelink/c3rDfqR8nn. All parsed data from wwPDB for the human membrane proteome are available at https://sid.erda.dk/sharelink/AtoGToVaZ8. All data related to the human membrane proteome and our X-ray subset are available at https://sid.erda.dk/sharelink/ds37GLjR8U. All data related to the MP stability benchmark are available at https://sid.erda.dk/sharelink/foHTKP49jC.

## SUPPLEMENTARY MATERIAL

An online supplement to this article can be found by visiting BJ Online at http://www.biophysj.org.

### Supplementary Figures

**Figure S1:**
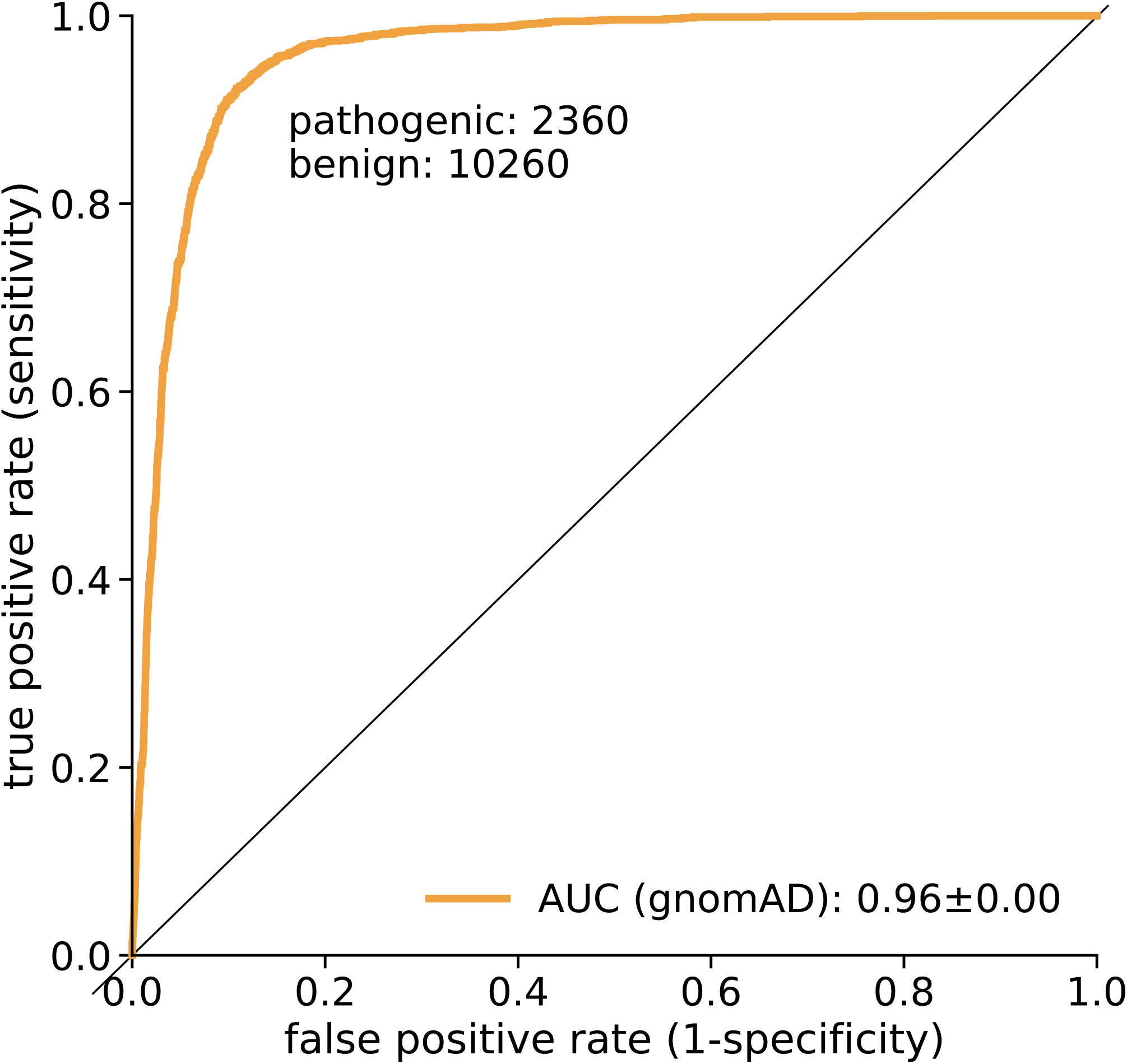
ROC analysis for gnomAD allele frequencies

#### Extended supplemental figure collection 1

**Neff values evaluated against GEMME coevolutionary score and sequence coverage per position:** `2022_11_11-ddE-neff-coverage.p` Per protein in the X-ray subset three plots are shown: first, the sequence coverage at each residue position vs. neff; second, the GEMME score (not normalized) vs. neff; third, the GEMME score vs. MSA sequence coverage. For the sequence coverage, our chosen threshold line is drawn at 50.

**Figure S2:**
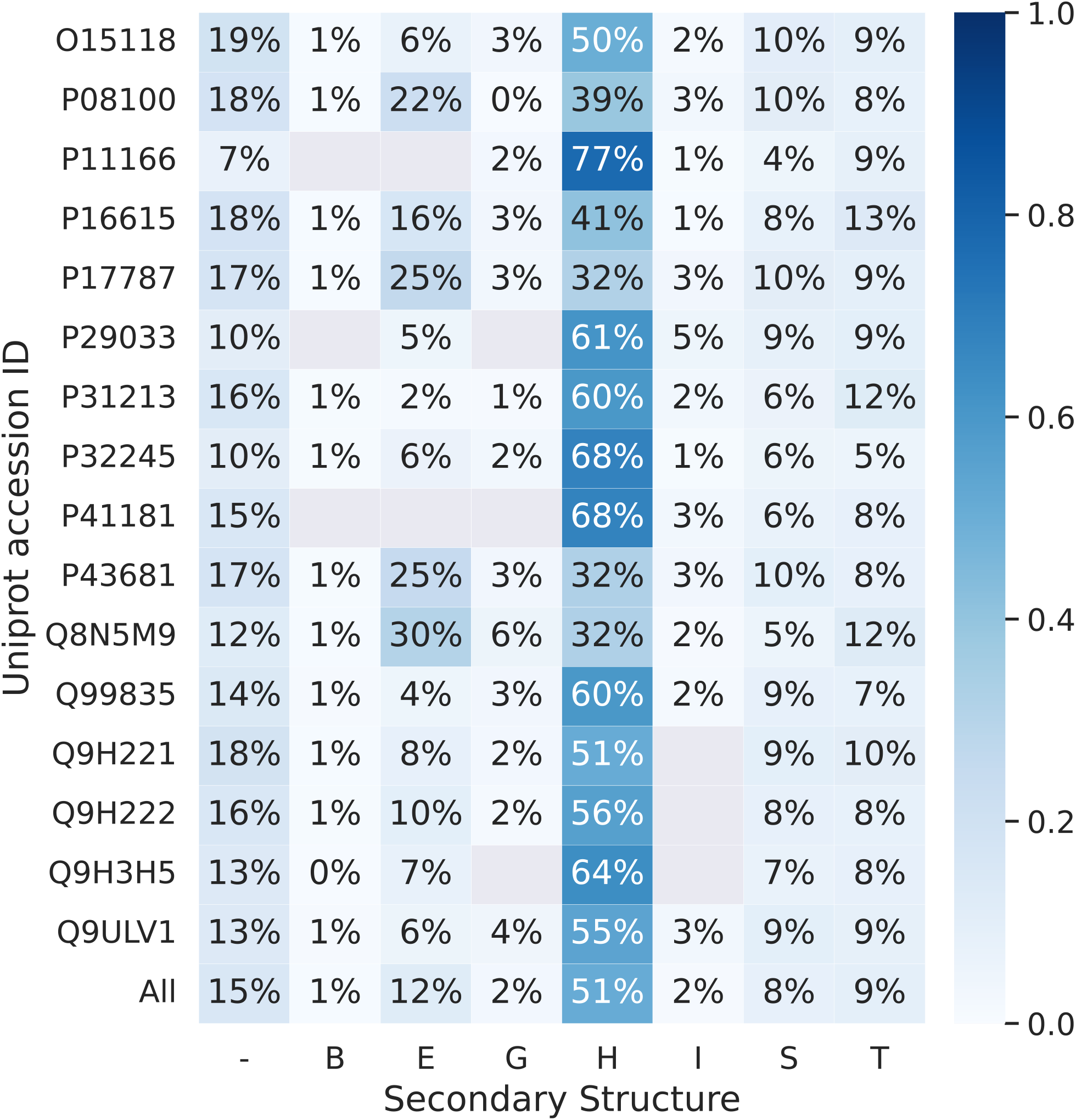
Secondary structure of target dataset calculated using DSSP (52). Abbreviation stand for H = α-helix; B = residue in isolated β-bridge; E = extended strand, participates in *β* ladder; G = 3-helix (3_10_ helix); I = 5 helix (π-helix); T = hydrogen bonded turn; S = bend; - = unstructured

**Figure S3:**
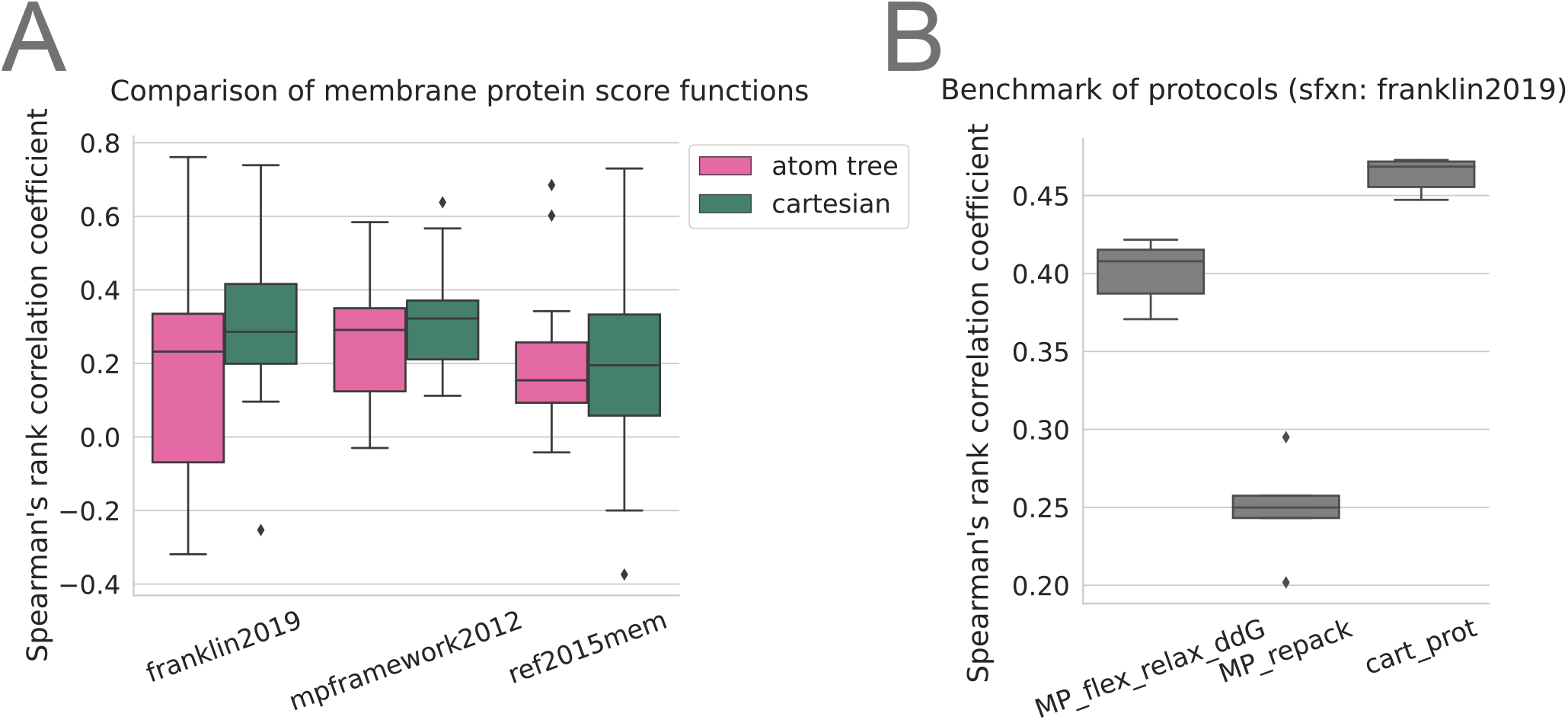
Benchmark of Rosetta ΔΔG calculations for MPs. (A) Comparison of accuracy of stability calculations performed with different membrane protein score functions but using the same protocol and data set. Data was extracted from (56). (B) Comparison using different protocols but the same score function (franklin2019; (35)) and was conducted on the benchmark set described in table S3.

**Figure S4:**
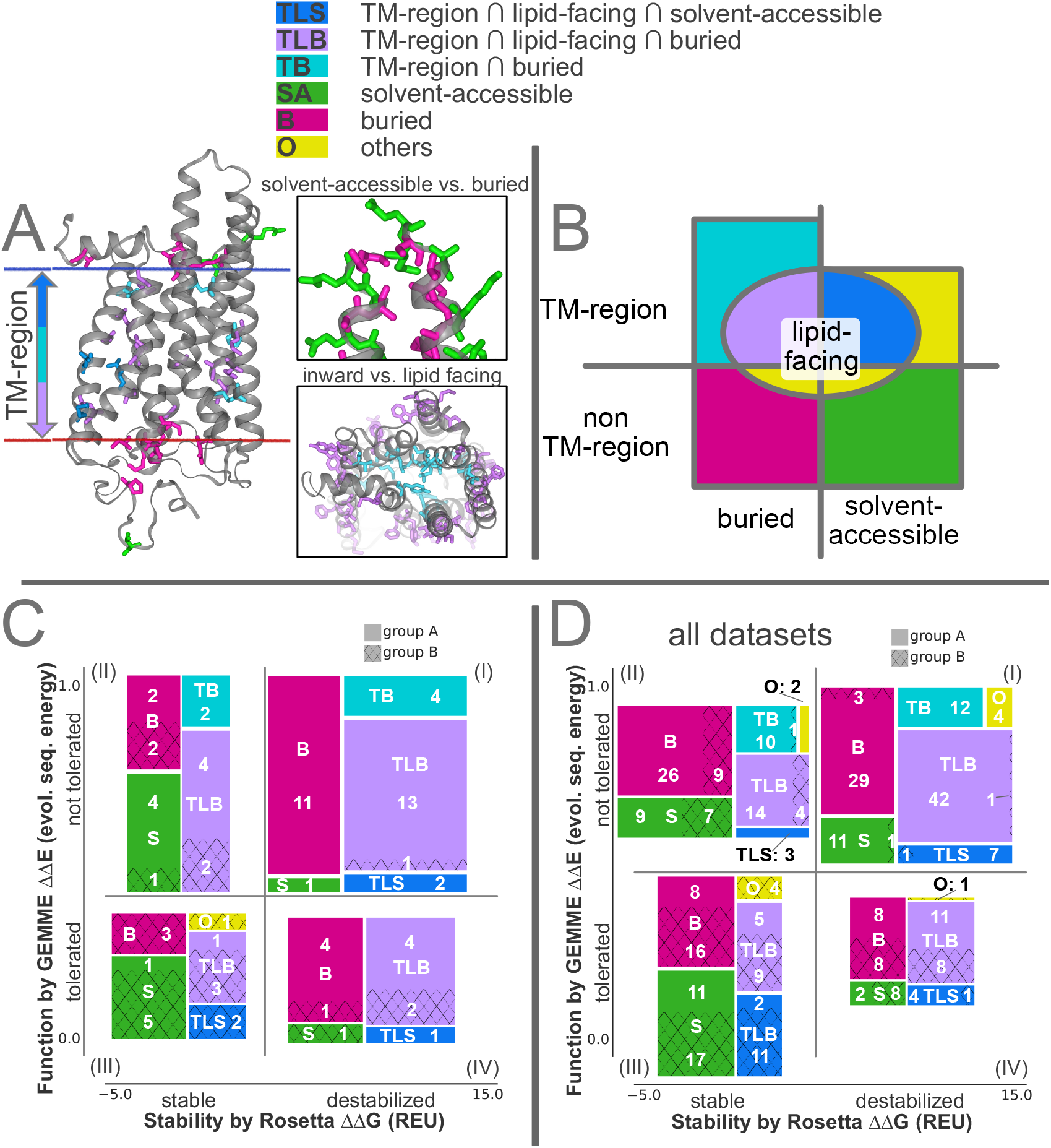
Protein regions and their overlaps and analysis of GPCR variant classification performance and structural differences of variant effects. (A) [left] Illustration of the different residue categories used in this work on OPSD, namely, whether they are inside (dark blue, turquoise, and violet) or outside of the transmembrane (TM) region (green and pink), and their orientation towards the membrane (lipid-facing: blue and violet), and whether they are solvent-accessible or buried. The structure on the left shows disease-associated variants, while the two inserts on the right illustrate solvent-accessible vs. buried, and inwards vs. lipid-facing, more generally and are not restricted to a disease relationship. Variants labeled with *other* are rare combinations, e.g., residues within the TM region that are solvent accessible but do not face the membrane, and some at the intersection between the membrane and the solvent (like lipid-facing but not placed within the TM region; see note in Limitations section). (B) Overview about the protein region distribution. (C) Similar as Fig. 3C, variant counts in the four quadrants, separated by their position in the proteins, are shown for *group A* (pathogenic, full) and *group B* (benign and/or non-rare, hashed) variants. (D) Variant counts as seen in (C) are shown summed over all 15 proteins.

### Supplementary Tables

#### Extended supplemental table 1

**Variant counts per protein:** `2022_11_11-count_hMP_anno_nonsyndel_PDB_publish.xlsx` The extended supplemental table is a collection of all displayed data and additional information on the data shown in the main manuscript and the supplement, including worksheet tabs:

- *2022_05_05-count_hMP_Clinvar_gnomad_PDB_nonsyndel_df*: contains a statistic of variant counts and PDB ids for each human membrane proteins that was experimentally resolved (data fetched by 2022-05-05)
- *Variant_annotation*: variant count from ClinVar and gnomAD for all human membrane proteins separated by cellular compartments
- *Variant_annotation_hP*: variant and protein count from ClinVar and gnomAD for all human proteins
- *Variant_annotation_PDB*: variant count from ClinVar and gnomAD for all human membrane proteins that are located in experimentally resolved protein regions, separated by cellular compartments
- *Category_variant_annotation*: variant count from ClinVar and gnomAD for all human membrane proteins separated by their membrane protein category and further subdivided into the variants located in the membrane bilayer; protein counts per category are also added.
- *Category_variant_annotation_PDB*: variant count from ClinVar and gnomAD for all human membrane proteins that are located in experimentally resolved protein regions, separated by their membrane protein category and further subdivided into the variants located in the membrane bilayer; protein counts per category are also added.
- *exp_ddg_benchmark*: data used for Rosetta stability benchmark (Supplementary Material table S3).
- *X-ray_set*: X-ray protein information table (equal to table 3
- *X-ray_set_app:* extended X-ray protein information table including variant counts after each sequential filtering steps and GEMME/MSA statistics
- *X-ray_set_app_AUC:* further extended X-ray protein information table (from worksheet tab *X-ray_set_app)* including additionally the AUC calculations (error via bootstrapping) for each filtered set of variants and the sequential filtered remaining data.
- *classes:* AUC and quadrant variant counts for each protein class in total and in the TM region.

#### Extended supplemental table 2

**Information of variants for each X-ray PDB with at least 1 benign and 1 pathogenic variant:** `2022_05_05-count_hMP_anno_splitPDB_Xi` The extended supplemental table is a collection of variant information (ClinVar and gnomAD per cellular compartment) per protein (each in a separate worksheet tab) per PDB-ID and chain. This is only generated for proteins, where at least one benign and one pathogenic variant is located in an experimentally resolved region. Further the StrucSel score (see Methods and Materials) is given, including further information about the PDB

**Table S1:** Number of variants annotated by ClinVar or gnomAD for those proteins with at least one experimentally resolved structure per cellular compartment

|  | all | Extracellular | Cytoplasmic | Transmembrane | other |
| --- | --- | --- | --- | --- | --- |
| total | 281,220 | 112,643 | 61,112 | 29,944 | 77,521 |
| benign | 2,089 | 957 | 495 | 169 | 4,029 |
| benign ∩ gnomAD | 1,951 | 883 | 467 | 155 | 446 |
| pathogenic | 4,030 | 1,430 | 1,118 | 807 | 3,822 |
| patho. ∩ gnomAD | 1,046 | 326 | 320 | 208 | 192 |
| VUS (ClinVar) | 18,882 | 7,592 | 5,962 | 1,974 | 3,354 |
| VUS ∩ gnomAD | 9,685 | 3,777 | 3,190 | 908 | 1,810 |
| only gnomAD | 256,219 | 102,664 | 53,537 | 26,994 | 73,024 |

**Table S2:** Variant counts, AUC and variant counts within the quadrants for each protein class. Cutoffs are taken from the complete dataset.

| protein class info |  |  | after filtering |  | AUC |  | Q I |  | Q II |  | Q III |  | Q IV |  |
| --- | --- | --- | --- | --- | --- | --- | --- | --- | --- | --- | --- | --- | --- | --- |
| class | # proteins | subselection | group A | group B | $\Delta\Delta G$ | $\Delta\Delta E$ | group A | group B | group A | group B | group A | group B | group A | group B |
| Cell Junction | 1 | all | 33 | 16 | 0.42 | 0.84 | 8 | 1 | 12 | 0 | 8 | 11 | 5 | 4 |
|  |  | TM region | 24 | 8 | 0.47 | 0.78 | 6 | 1 | 9 | 0 | 4 | 6 | 5 | 1 |
| Enzyme | 2 | all | 24 | 10 | 0.62 | 0.83 | 12 | 0 | 8 | 3 | 2 | 4 | 2 | 3 |
|  |  | TM region | 6 | 33 | 1 | 0.83 | 4 | 0 | 0 | 1 | 0 | 1 | 2 | 1 |
| GPCR | 4 | all | 54 | 24 | 0.79 | 0.83 | 31 | 1 | 12 | 5 | 2 | 14 | 9 | 4 |
|  |  | TM region | 31 | 11 | 0.81 | 0.87 | 19 | 1 | 6 | 2 | 1 | 5 | 5 | 3 |
| Ion channel | 3 | all | 11 | 12 | 0.72 | 0.81 | 6 | 1 | 3 | 3 | 2 | 8 | 0 | 20 |
|  |  | TM region | 4 | 3 | 0.42 | 0.75 | 1 | 0 | 2 | 1 | 1 | 2 | 0 | 0 |
| Transporter | 5 | all | 98 | 42 | 0.63 | 0.82 | 48 | 3 | 29 | 9 | 12 | 20 | 9 | 10 |
|  |  | TM region | 44 | 10 | 0.71 | 0.97 | 27 | 0 | 11 | 1 | 2 | 6 | 4 | 3 |

**Table S3:**
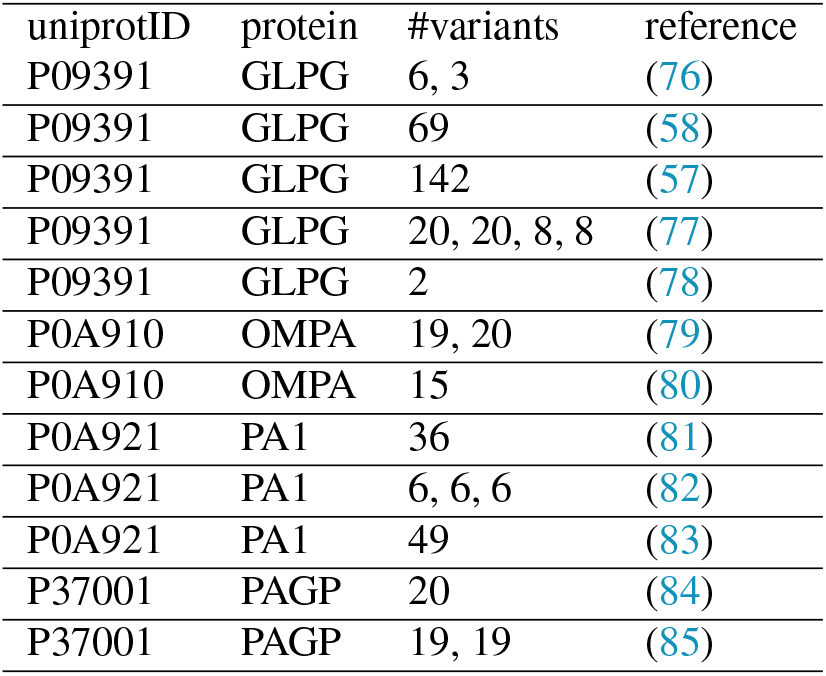
Experimental ΔΔG datasets (all from E. coli) used for benchmarking. Multiple variant counts indicate different pH, labeling tags or temperatures.

## REFERENCES

1. Soskine, M., and D. S. Tawfik, 2010. Mutational effects and the evolution of new protein functions. Nature Reviews Genetics 11:572–582. https://www.nature.com/articles/nrg2808.

2. Pey, A. L., F. Stricher, L. Serrano, and A. Martinez, 2007. Predicted effects of missense mutations on native-state stability account for phenotypic outcome in phenylketonuria, a paradigm of misfolding diseases. American Journal of Human Genetics 81:1006–1024.

3. Yue, P., Z. Li, and J. Moult, 2005. Loss of protein structure stability as a major causative factor in monogenic disease. Journal of Molecular Biology 353:459–473.

4. Casadio, R., M. Vassura, S. Tiwari, P. Fariselli, and P. Luigi Martelli, 2011. Correlating disease-related mutations to their effect on protein stability: a large-scale analysis of the human proteome. Human Mutation 32:1161–1170.

5. Martelli, P. L., P. Fariselli, C. Savojardo, G. Babbi, F. Aggazio, and R. Casadio, 2016. Large scale analysis of protein stability in OMIM disease related human protein variants. BMC genomics 17 Suppl 2:397.

6. Nielsen, S. V., A. Stein, A. B. Dinitzen, E. Papaleo, M. H. Tatham, E. G. Poulsen, M. M. Kassem, L. J. Rasmussen, K. Lindorff-Larsen, and R. Hartmann-Petersen, 2017. Predicting the impact of Lynch syndrome-causing missense mutations from structural calculations. PLoS genetics 13:e1006739.

7. Abildgaard, A. B., A. Stein, S. V. Nielsen, K. Schultz-Knudsen, E. Papaleo, A. Shrikhande, E. R. Hoffmann, I. Bernstein, A.-M. Gerdes, M. Takahashi, C. Ishioka, K. Lindorff-Larsen, and R. Hartmann-Petersen, 2019. Computational and cellular studies reveal structural destabilization and degradation of MLH1 variants in Lynch syndrome. eLife 8:e49138.

8. Gersing, S. K., Y. Wang, M. Grønbæk-Thygesen, C. Kampmeyer, L. Clausen, M. Willemoës, C. Andréasson, A. Stein, K. Lindorff-Larsen, and R. Hartmann-Petersen, 2021. Mapping the degradation pathway of a disease-linked aspartoacylase variant. PLoS genetics 17:e1009539.

9. Scheller, R., A. Stein, S. V. Nielsen, F. I. Marin, A.-M. Gerdes, M. Di Marco, E. Papaleo, K. Lindorff-Larsen, and R. Hartmann-Petersen, 2019. Toward mechanistic models for genotype-phenotype correlations in phenylketonuria using protein stability calculations. Human Mutation 40:444–457.

10. Clausen, L., A. Stein, M. Grønbæk-Thygesen, L. Nygaard, C. L. Søltoft, S. V. Nielsen, M. Lisby, T. Ravid, K. Lindorff-Larsen, and R. Hartmann-Petersen, 2020. Folliculin variants linked to Birt-Hogg-Dubé syndrome are targeted for proteasomal degradation. PLoS genetics 16:e1009187.

11. Stein, A., D. M. Fowler, R. Hartmann-Petersen, and K. Lindorff-Larsen, 2019. Biophysical and Mechanistic Models for Disease-Causing Protein Variants. Trends in Biochemical Sciences 44:575–588. http://www.sciencedirect.com/science/article/pii/S0968000419300039.

12. Park, H., P. Bradley, P. Greisen, Y. Liu, V. K. Mulligan, D. E. Kim, D. Baker, and F. DiMaio, 2016. Simultaneous Optimization of Biomolecular Energy Functions on Features from Small Molecules and Macromolecules. Journal of Chemical Theory and Computation 12:6201–6212.

13. Guerois, R., J. E. Nielsen, and L. Serrano, 2002. Predicting changes in the stability of proteins and protein complexes: a study of more than 1000 mutations. Journal of Molecular Biology 320:369–387.

14. Kellogg, E. H., A. Leaver-Fay, and D. Baker, 2011. Role of conformational sampling in computing mutation-induced changes in protein structure and stability. Proteins 79:830–838.

15. Ó Conchúir, S., K. A. Barlow, R. A. Pache, N. Ollikainen, K. Kundert, M. J. O’Meara, C. A. Smith, and T. Kortemme, 2015. A Web Resource for Standardized Benchmark Datasets, Metrics, and Rosetta Protocols for Macromolecular Modeling and Design. PloS One 10:e0130433.

16. Frenz, B., S. M. Lewis, I. King, F. DiMaio, H. Park, and Y. Song, 2020. Prediction of Protein Mutational Free Energy: Benchmark and Sampling Improvements Increase Classification Accuracy. Frontiers in Bioengineering and Biotechnology 8:558247.

17. Jepsen, M. M., D. M. Fowler, R. Hartmann-Petersen, A. Stein, and K. Lindorff-Larsen, 2020. Chapter 5 - Classifying disease-associated variants using measures of protein activity and stability. In A. L. Pey, editor, Protein Homeostasis Diseases, Academic Press, 91–107. https://www.sciencedirect.com/science/article/pii/B9780128191323000051.

18. Cagiada, M., K. E. Johansson, A. Valanciute, S. V. Nielsen, R. Hartmann-Petersen, J. J. Yang, D. M. Fowler, A. Stein, and K. Lindorff-Larsen, 2021. Understanding the Origins of Loss of Protein Function by Analyzing the Effects of Thousands of Variants on Activity and Abundance. Molecular Biology and Evolution 38:3235–3246.

19. Høie, M. H., M. Cagiada, A. H. Beck Frederiksen, A. Stein, and K. Lindorff-Larsen, 2022. Predicting and interpreting large-scale mutagenesis data using analyses of protein stability and conservation. Cell Reports 38:110207.

20. Meng, X., J. Clews, V. Kargas, X. Wang, and R. C. Ford, 2017. The cystic fibrosis transmembrane conductance regulator (CFTR) and its stability. Cellular and molecular life sciences: CMLS 74:23–38.

21. Kampmeyer, C., S. V. Nielsen, L. Clausen, A. Stein, A.-M. Gerdes, K. Lindorff-Larsen, and R. Hartmann-Petersen, 2017. Blocking protein quality control to counter hereditary cancers. Genes, Chromosomes & Cancer 56:823–831.

22. Uhlén, M., L. Fagerberg, B. M. Hallström, C. Lindskog, P. Oksvold, A. Mardinoglu, A. Sivertsson, C. Kampf, E. Sjöstedt, A. Asplund, I. Olsson, K. Edlund, E. Lundberg, S. Navani, C. A.-K. Szigyarto, J. Odeberg, D. Djureinovic, J. O. Takanen, S. Hober, T. Alm, P.-H. Edqvist, H. Berling, H. Tegel, J. Mulder, J. Rockberg, P. Nilsson, J. M. Schwenk, M. Hamsten, K. von Feilitzen, M. Forsberg, L. Persson, F. Johansson, M. Zwahlen, G. von Heijne, J. Nielsen, and F. Pontén, 2015. Proteomics. Tissue-based map of the human proteome. Science (New York, N.Y.) 347:1260419.

23. von Heijne, G., 2007. The membrane protein universe: what’s out there and why bother? Journal of Internal Medicine 261:543–557.

24. Hauser, A. S., M. M. Attwood, M. Rask-Andersen, H. B. Schiöth, and D. E. Gloriam, 2017. Trends in GPCR drug discovery: new agents, targets and indications. Nature Reviews Drug Discovery 16:829–842. https://www.nature.com/articles/nrd.2017.178.

25. Sanders, C. R., and J. K. Nagy, 2000. Misfolding of membrane proteins in health and disease: the lady or the tiger? Current Opinion in Structural Biology 10:438–442.

26. Hamel, C., 2006. Retinitis pigmentosa. Orphanet Journal of Rare Diseases 1:40.

27. Koepsell, H., 2020. Glucose transporters in brain in health and disease. Pflugers Archiv: European Journal of Physiology 472:1299–1343.

28. Vanier, M. T., 2010. Niemann-Pick disease type C. Orphanet Journal of Rare Diseases 5:16.

29. Cournia, Z., T. W. Allen, I. Andricioaei, B. Antonny, D. Baum, G. Brannigan, N.-V. Buchete, J. T. Deckman, L. Delemotte, C. del Val, R. Friedman, P. Gkeka, H.-C. Hege, J. Hénin, M. A. Kasimova, A. Kolocouris, M. L. Klein, S. Khalid, M. J. Lemieux, N. Lindow, M. Roy, J. Selent, M. Tarek, F. Tofoleanu, S. Vanni, S. Urban, D. J. Wales, J. C. Smith, and A.-N. Bondar, 2015. Membrane Protein Structure, Function and Dynamics: A Perspective from Experiments and Theory. The Journal of membrane biology 248:611–640. https://www.ncbi.nlm.nih.gov/pmc/articles/PMC4515176/.

30. Hong, H., 2015. Role of Lipids in Folding, Misfolding and Function of Integral Membrane Proteins. In O. Gursky, editor, Lipids in Protein Misfolding, Springer International Publishing, Cham, Advances in Experimental Medicine and Biology, 1–31. https://doi.org/10.1007/978-3-319-17344-3_1.

31. Booth, P. J., and J. Clarke, 2010. Membrane protein folding makes the transition. Proceedings of the National Academy of Sciences 107:3947–3948. https://www.pnas.org/content/107/9/3947.

32. Chang, Y.-C., and J. U. Bowie, 2014. Measuring membrane protein stability under native conditions. Proceedings of the National Academy of Sciences 111:219–224. https://www.pnas.org/doi/full/10.1073/pnas.1318576111, publisher: Proceedings of the National Academy of Sciences.

33. Boland, C., S. Olatunji, J. Bailey, N. Howe, D. Weichert, S. K. Fetics, X. Yu, J. Merino-Gracia, C. Delsaut, and M. Caffrey, 2018. Membrane (and Soluble) Protein Stability and Binding Measurements in the Lipid Cubic Phase Using Label-Free Differential Scanning Fluorimetry. Analytical Chemistry 90:12152–12160. https://doi.org/10.1021/acs.analchem.8b03176, publisher: American Chemical Society.

34. Marx, D. C., and K. G. Fleming, 2021. Membrane proteins enter the fold. Current Opinion in Structural Biology 69:124–130. https://www.sciencedirect.com/science/article/pii/S0959440X21000440.

35. Alford, R. F., P. J. Fleming, K. G. Fleming, and J. J. Gray, 2021. Protein Structure Prediction and Design in a Biologically Realistic Implicit Membrane. Biophysical Journal 120:4635. https://www.sciencedirect.com/science/article/pii/S0006349521007530.

36. Laine, E., Y. Karami, and A. Carbone, 2019. GEMME: a simple and fast global epistatic model predicting mutational effects. Molecular Biology and Evolution msz179.

37. Frazer, J., P. Notin, M. Dias, A. Gomez, J. K. Min, K. Brock, Y. Gal, and D. S. Marks, 2021. Disease variant prediction with deep generative models of evolutionary data. Nature 599:91–95.

38. Feinauer, C., and M. Weigt, 2017. Context-Aware Prediction of Pathogenicity of Missense Mutations Involved in Human Disease. Technical report, bioRxiv. https://www.biorxiv.org/content/10.1101/103051v1.

39. Nicoludis, J. M., and R. Gaudet, 2018. Applications of sequence coevolution in membrane protein biochemistry. Biochimica et Biophysica Acta (BBA) - Biomembranes 1860:895–908. https://www.sciencedirect.com/science/article/pii/S0005273617303140.

40. Lin, Z., H. Akin, R. Rao, B. Hie, Z. Zhu, W. Lu, N. Smetanin, R. Verkuil, O. Kabeli, Y. Shmueli, A. d. S. Costa, M. Fazel-Zarandi, T. Sercu, S. Candido, and A. Rives, 2022. Evolutionary-scale prediction of atomic level protein structure with a language model. https://www.biorxiv.org/content/10.1101/2022.07.20.500902v2, pages: 2022.07.20.500902 Section: New Results.

41. Gerasimavicius, L., B. J. Livesey, and J. A. Marsh, 2022. Loss-of-function, gain-of-function and dominant-negative mutations have profoundly different effects on protein structure: implications for variant effect prediction. https://www.biorxiv.org/content/10.1101/2021.10.23.465554v2, pages: 2021.10.23.465554 Section: New Results.

42. The UniProt Consortium, 2021. UniProt: the universal protein knowledgebase in 2021. Nucleic Acids Research 49:D480–D489. https://doi.org/10.1093/nar/gkaa1100.

43. Landrum, M. J., J. M. Lee, M. Benson, G. R. Brown, C. Chao, S. Chitipiralla, B. Gu, J. Hart, D. Hoffman, W. Jang, K. Karapetyan, K. Katz, C. Liu, Z. Maddipatla, A. Malheiro, K. McDaniel, M. Ovetsky, G. Riley, G. Zhou, J. B. Holmes, B. L. Kattman, and D. R. Maglott, 2018. ClinVar: improving access to variant interpretations and supporting evidence. Nucleic Acids Research 46:D1062–D1067.

44. Karczewski, K. J., L. C. Francioli, G. Tiao, B. B. Cummings, J. Alföldi, Q. Wang, R. L. Collins, K. M. Laricchia, A. Ganna, D. P. Birnbaum, L. D. Gauthier, H. Brand, M. Solomonson, N. A. Watts, D. Rhodes, M. Singer-Berk, E. M. England, E. G. Seaby, J. A. Kosmicki, R. K. Walters, K. Tashman, Y. Farjoun, E. Banks, T. Poterba, A. Wang, C. Seed, N. Whiffin, J. X. Chong, K. E. Samocha, E. Pierce-Hoffman, Z. Zappala, A. H. O’Donnell-Luria, E. V. Minikel, B. Weisburd, M. Lek, J. S. Ware, C. Vittal, I. M. Armean, L. Bergelson, K. Cibulskis, K. M. Connolly, M. Covarrubias, S. Donnelly, S. Ferriera, S. Gabriel, J. Gentry, N. Gupta, T. Jeandet, D. Kaplan, C. Llanwarne, R. Munshi, S. Novod, N. Petrillo, D. Roazen, V. Ruano-Rubio, A. Saltzman, M. Schleicher, J. Soto, K. Tibbetts, C. Tolonen, G. Wade, M. E. Talkowski, B. M. Neale, M. J. Daly, and D. G. MacArthur, 2020. The mutational constraint spectrum quantified from variation in 141,456 humans. Nature 581:434–443. https://www.nature.com/articles/s41586-020-2308-7.

45. Cock, P. J., T. Antao, J. T. Chang, B. A. Chapman, C. J. Cox, A. Dalke, I. Friedberg, T. Hamelryck, F. Kauff, B. Wilczynski, et al., 2009. Biopython: freely available Python tools for computational molecular biology and bioinformatics. Bioinformatics 25:1422–1423.

46. Remmert, M., A. Biegert, A. Hauser, and J. Söding, 2011. HHblits: lightning-fast iterative protein sequence searching by HMM-HMM alignment. Nature Methods 9:173–175.

47. Mirdita, M., L. von den Driesch, C. Galiez, M. J. Martin, J. Söding, and M. Steinegger, 2017. Uniclust databases of clustered and deeply annotated protein sequences and alignments. Nucleic Acids Research 45:D170–D176.

48. UniProt Consortium, 2019. UniProt: a worldwide hub of protein knowledge. Nucleic Acids Research 47:D506–D515.

49. Ruan, J., Z. Liu, M. Sun, Y. Wang, J. Yue, and G. Yu, 2019. DBS: a fast and informative segmentation algorithm for DNA copy number analysis. BMC bioinformatics 20:1.

50. Koehler Leman, J., S. Lyskov, and R. Bonneau, 2017. Computing structure-based lipid accessibility of membrane proteins with mp_lipid_acc in RosettaMP. BMC Bioinformatics 18:115. https://doi.org/10.1186/s12859-017-1541-z.

51. Kabsch, W., and C. Sander, 1983. Dictionary of protein secondary structure: pattern recognition of hydrogen-bonded and geometrical features. Biopolymers 22:2577–2637.

52. Touw, W. G., C. Baakman, J. Black, T A. H. te Beek, E. Krieger, R. P. Joosten, and G. Vriend, 2015. A series of PDB-related databanks for everyday needs. Nucleic Acids Research 43:D364–368.

53. Lomize, M. A., I. D. Pogozheva, H. Joo, H. I. Mosberg, and A. L. Lomize, 2012. OPM database and PPM web server: resources for positioning of proteins in membranes. Nucleic Acids Research 40:D370–376.

54. Alford, R. F., J. K. Leman, B. D. Weitzner, A. M. Duran, D. C. Tilley, A. Elazar, and J. J. Gray, 2015. An Integrated Framework Advancing Membrane Protein Modeling and Design. PLOS Computational Biology 11:e1004398. https://journals.plos.org/ploscompbiol/article?id=10.1371/journal.pcbi.1004398.

55. Koehler Leman, J., B. K. Mueller, and J. J. Gray, 2017. Expanding the toolkit for membrane protein modeling in Rosetta. Bioinformatics 33:754–756. https://doi.org/10.1093/bioinformatics/btw716.

56. Koehler Leman, J., S. Lyskov, S. M. Lewis, J. Adolf-Bryfogle, R. F. Alford, K. Barlow, Z. Ben-Aharon, D. Farrell, J. Fell, W. A. Hansen, A. Harmalkar, J. Jeliazkov, G. Kuenze, J. D. Krys, A. Ljubetič, A. L. Loshbaugh, J. Maguire, R. Moretti, V. K. Mulligan, M. L. Nance, P. T. Nguyen, S. Ó Conchúir, S. S. Roy Burman, R. Samanta, S. T. Smith, F. Teets, J. K. S. Tiemann, A. Watkins, H. Woods, B. J. Yachnin, C. D. Bahl, C. Bailey-Kellogg, D. Baker, R. Das, F. DiMaio, S. D. Khare, T. Kortemme, J. W. Labonte, K. Lindorff-Larsen, J. Meiler, W. Schief, O. Schueler-Furman, J. B. Siegel, A. Stein, V. Yarov-Yarovoy, B. Kuhlman, A. Leaver-Fay, D. Gront, J. J. Gray, and R. Bonneau, 2021. Ensuring scientific reproducibility in bio-macromolecular modeling via extensive, automated benchmarks. Nature Communications 12:6947. https://www.nature.com/articles/s41467-021-27222-7.

57. Baker, R. P., and S. Urban, 2012. Architectural and thermodynamic principles underlying intramembrane protease function. Nature Chemical Biology 8:759–768. https://www.nature.com/articles/nchembio.1021.

58. Paslawski, W., O. K. Lillelund, J. V. Kristensen, N. P. Schafer, R. P. Baker, S. Urban, and D. E. Otzen, 2015. Cooperative folding of a polytopic α-helical membrane protein involves a compact N-terminal nucleus and nonnative loops. Proceedings of the National Academy of Sciences 112:7978–7983. https://www.pnas.org/content/112/26/7978.

59. Krzanowski, W. J., and D. J. Hand, 2009. ROC Curves for Continuous Data. Chapman and Hall/CRC, New York.

60. Fleishman, S. J., A. Leaver-Fay, J. E. Corn, E.-M. Strauch, S. D. Khare, N. Koga, J. Ashworth, P. Murphy, F. Richter, G. Lemmon, J. Meiler, and D. Baker, 2011. RosettaScripts: A Scripting Language Interface to the Rosetta Macromolecular Modeling Suite. PLOS ONE 6:e20161. https://journals.plos.org/plosone/article?id=10.1371/journal.pone.0020161, publisher: Public Library of Science.

61. Khatib, F., S. Cooper, M. D. Tyka, K. Xu, I. Makedon, Z. Popović, D. Baker, and F. Players, 2011. Algorithm discovery by protein folding game players. Proceedings of the National Academy of Sciences 108:18949–18953. https://www.pnas.org/doi/full/10.1073/pnas.1115898108, publisher: Proceedings of the National Academy of Sciences.

62. Maguire, J. B., H. K. Haddox, D. Strickland, S. F. Halabiya, B. Coventry, J. R. Griffin, S. V. S. R. K. Pulavarti, M. Cummins, D. F. Thieker, E. Klavins, T. Szyperski, F. DiMaio, D. Baker, and B. Kuhlman, 2021. Perturbing the energy landscape for improved packing during computational protein design. Proteins: Structure, Function, and Bioinformatics 89:436–449. https://onlinelibrary.wiley.com/doi/abs/10.1002/prot.26030, _eprint: https://onlinelibrary.wiley.com/doi/pdf/10.1002/prot.26030.

63. Zaucha, J., M. Heinzinger, A. Kulandaisamy, E. Kataka, a. L. Salvádor, P Popov, B. Rost, M. M. Gromiha, B. S. Zhorov, and D. Frishman, 2021. Mutations in transmembrane proteins: diseases, evolutionary insights, prediction and comparison with globular proteins. Briefings in Bioinformatics 22:bbaa132. https://doi.org/10.1093/bib/bbaa132.

64. Lee, E., and C. Manoil, 1994. Mutations eliminating the protein export function of a membrane-spanning sequence. Journal of Biological Chemistry 269:28822–28828. https://www.sciencedirect.com/science/article/pii/S0021925819619800.

65. Varadi, M., S. Anyango, M. Deshpande, S. Nair, C. Natassia, G. Yordanova, D. Yuan, O. Stroe, G. Wood, A. Laydon, A. Žídek, T. Green, K. Tunyasuvunakool, S. Petersen, J. Jumper, E. Clancy, R. Green, A. Vora, M. Lutfi, M. Figurnov, A. Cowie, N. Hobbs, P. Kohli, G. Kleywegt, E. Birney, D. Hassabis, and S. Velankar, 2021. AlphaFold Protein Structure Database: massively expanding the structural coverage of protein-sequence space with high-accuracy models. Nucleic Acids Research gkab1061. https://doi.org/10.1093/nar/gkab1061.

66. del Alamo, D., D. Sala, H. S. Mchaourab, and J. Meiler, 2022. Sampling alternative conformational states of transporters and receptors with AlphaFold2. eLife 11:e75751. https://doi.org/10.7554/eLife.75751.

67. Akdel, M., D. E. V. Pires, E. P. Pardo, J. Jänes, A. O. Zalevsky, B. Mészáros, P. Bryant, L. L. Good, R. A. Laskowski, G. Pozzati, A. Shenoy, W. Zhu, P. Kundrotas, V. R. Serra, C. H. M. Rodrigues, A. S. Dunham, D. Burke, N. Borkakoti, S. Velankar, A. Frost, K. Lindorff-Larsen, A. Valencia, S. Ovchinnikov, J. Durairaj, D. B. Ascher, J. M. Thornton, N. E. Davey, A. Stein, A. Elofsson, T. I. Croll, and P. Beltrao, 2021. A structural biology community assessment of AlphaFold 2 applications. Technical report, bioRxiv. https://www.biorxiv.org/content/10.1101/2021.09.26.461876v1.

68. wwPDB consortium, 2019. Protein Data Bank: the single global archive for 3D macromolecular structure data. Nucleic Acids Research 47:D520–D528. https://doi.org/10.1093/nar/gky949.

69. Sörmann, J., M. Schewe, P. Proks, T. Jouen-Tachoire, S. Rao, E. B. Riel, K. E. Agre, A. Begtrup, J. Dean, M. Descartes, J. Fischer, A. Gardham, C. Lahner, P. R. Mark, S. Muppidi, P. N. Pichurin, J. Porrmann, J. Schallner, K. Smith, V. Straub, P. Vasudevan, R. Willaert, E. P. Carpenter, K. E. J. Rödström, M. G. Hahn, T. Müller, T. Baukrowitz, M. E. Hurles, C. F. Wright, and S. J. Tucker, 2022. Gain-of-function mutations in KCNK3 cause a developmental disorder with sleep apnea. Nature Genetics 54:1534–1543. https://www.nature.com/articles/s41588-022-01185-x, number: 10 Publisher: Nature Publishing Group.

70. Custódio, T. F., P A. Paulsen, K. M. Frain, and B. P. Pedersen, 2021. Structural comparison of GLUT1 to GLUT3 reveal transport regulation mechanism in sugar porter family. Life Science Alliance 4. https://www.life-science-alliance.org/content/4/4/e202000858.

71. Hofmann, K. P., P. Scheerer, P W. Hildebrand, H.-W. Choe, J. H. Park, M. Heck, and O. P. Ernst, 2009. A G protein-coupled receptor at work: the rhodopsin model. Trends in Biochemical Sciences 34:540–552.

72. Kapoor, K., J. S. Finer-Moore, B. P. Pedersen, L. Caboni, A. Waight, R. C. Hillig, P. Bringmann, I. Heisler, T. Müller, H. Siebeneicher, and R. M. Stroud, 2016. Mechanism of inhibition of human glucose transporter GLUT1 is conserved between cytochalasin B and phenylalanine amides. Proceedings of the National Academy of Sciences of the United States of America 113:4711–4716. https://europepmc.org/articles/PMC4855560.

73. Zhao, G., and E. London, 2006. An amino acid “transmembrane tendency” scale that approaches the theoretical limit to accuracy for prediction of transmembrane helices: relationship to biological hydrophobicity. Protein Science: A Publication of the Protein Society 15:1987–2001.

74. Anderson, C. L., S. Munawar, L. Reilly, T. J. Kamp, C. T. January, B. P. Delisle, and L. L. Eckhardt, 2022. How Functional Genomics Can Keep Pace With VUS Identification. Frontiers in Cardiovascular Medicine 9. https://www.frontiersin.org/articles/10.3389/fcvm.2022.900431.

75. Jumper, J., R. Evans, A. Pritzel, T. Green, M. Figurnov, O. Ronneberger, K. Tunyasuvunakool, R. Bates, A. Žídek, A. Potapenko, A. Bridgland, C. Meyer, S. A. A. Kohl, A. J. Ballard, A. Cowie, B. Romera-Paredes, S. Nikolov, R. Jain, J. Adler, T. Back, S. Petersen, D. Reiman, E. Clancy, M. Zielinski, M. Steinegger, M. Pacholska, T. Berghammer, S. Bodenstein, D. Silver, O. Vinyals, A. W. Senior, K. Kavukcuoglu, P. Kohli, and D. Hassabis, 2021. Highly accurate protein structure prediction with AlphaFold. Nature 596:583–589. https://www.nature.com/articles/s41586-021-03819-2, number: 7873 Publisher: Nature Publishing Group.

76. Gaffney, K. A., and H. Hong, 2018. The rhomboid protease GlpG has weak interaction energies in its active site hydrogen bond network. Journal of General Physiology 151:282–291. https://doi.org/10.1085/jgp.201812047.

77. Guo, R., K. Gaffney, Z. Yang, M. Kim, S. Sungsuwan, X. Huang, W. L. Hubbell, and H. Hong, 2016. Steric trapping reveals a cooperativity network in the intramembrane protease GlpG. Nature Chemical Biology 12:353–360. https://www.nature.com/articles/nchembio.2048.

78. Min, D., R. E. Jefferson, J. U. Bowie, and T.-Y. Yoon, 2015. Mapping the energy landscape for second-stage folding of a single membrane protein. Nature Chemical Biology 11:981–987. https://www.nature.com/articles/nchembio.1939.

79. Hong, H., S. Park, R. H. Flores Jiménez, D. Rinehart, and L. K. Tamm, 2007. Role of Aromatic Side Chains in the Folding and Thermodynamic Stability of Integral Membrane Proteins. Journal of the American Chemical Society 129:8320–8327. https://doi.org/10.1021/ja068849o.

80. Hong, H., G. Szabo, and L. K. Tamm, 2006. Electrostatic couplings in OmpA ion-channel gating suggest a mechanism for pore opening. Nature Chemical Biology 2:627–635. https://www.nature.com/articles/nchembio827.

81. Moon, C. P., and K. G. Fleming, 2011. Side-chain hydrophobicity scale derived from transmembrane protein folding into lipid bilayers. Proceedings of the National Academy of Sciences 108:10174–10177. https://www.pnas.org/content/108/25/10174.

82. Stanley, A. M., and K. G. Fleming, 2007. The Role of a Hydrogen Bonding Network in the Transmembrane jß-Barrel OMPLA. Journal of Molecular Biology 370:912–924. https://www.sciencedirect.com/science/article/pii/S0022283607006213.

83. McDonald, S. K., and K. G. Fleming, 2016. Aromatic Side Chain Water-to-Lipid Transfer Free Energies Show a Depth Dependence across the Membrane Normal. Journal of the American Chemical Society 138:7946–7950. https://doi.org/10.1021/jacs.6b03460.

84. Marx, D. C., and K. G. Fleming, 2017. Influence of Protein Scaffold on Side-Chain Transfer Free Energies. Biophysical Journal 113:597–604. https://www.cell.com/biophysj/abstract/S0006-3495(17)30682-3.

85. Huysmans, G. H. M., S. A. Baldwin, D. J. Brockwell, and S. E. Radford, 2010. The transition state for folding of an outer membrane protein. Proceedings of the National Academy of Sciences 107:4099–4104. https://www.pnas.org/content/107/9/4099.

